# IL-1α drives a tumor-stroma-neutrophil axis through inflammatory fibroblast activation in head and neck cancer

**DOI:** 10.64898/2026.01.20.700440

**Authors:** Eva-Maria Hanschmann, Kirsten Bruderek, Francisca Hofmann-Vega, Mathias Schmidt, Bettina Budeus, Wibke Oberlies, Xiang Wei, Mathis Richter, Simon Lemm, Jagoda Agnieszka Szlachetko, Sarah Ryan, Almke Bader, Anke Daser, Stephan Lang, Daniela Maier-Begandt, Timon Hussain, Oliver Söhnlein, Sven Brandau

## Abstract

In head and neck squamous cell carcinoma (HNSCC), high tumor-associated neutrophil (TAN) density is a robust biomarker of poor prognosis. TANs predominantly localize to the stroma and their intratumoral density strongly correlates with adverse outcome. Here, we investigated how tumor-stroma communication regulates TAN recruitment, activation, and spatial organization.

We identified interleukin-1α (IL-1α), released by viable or necrotic tumor cells, as an upstream signal that induces an inflammatory cancer associated fibroblast (iCAF)-like transcriptional program in patient-derived mesenchymal stromal cells (MSCs) of the oral cavity. IL-1α-stimulated MSCs secrete factors associated with neutrophil recruitment, survival, and function, together with mediators of extracellular matrix remodeling and angiogenesis. Notably, the IL-1α-induced iCAF-like transcriptional program closely resembles CAF subsets in HNSCC that are associated with poor clinical outcome.

Functionally, conditioned media from IL-1α-stimulated MSCs promoted tumor growth and enhanced polymorphonuclear neutrophil survival, activation, trans-well migration and infiltration into spheroids, *in vitro*. In zebrafish xenografts, co-injection of IL-1α-overexpressing tumor cells and MSCs markedly amplified neutrophil infiltration.

TCGA analysis demonstrated robust correlations between the IL-1α-induced MSC gene signature and neutrophil signatures across multiple TAN subsets in human HNSCC. Spatial analysis of HNSCC tissues showed that stromal regions adjacent to *IL1A*-positive tumor islets were enriched for *CXCL8/CSF3* double-positive cells and exhibited increased TAN density, including higher frequencies of NE– and MPO-positive neutrophils.

Collectively, these findings define an IL-1α-dependent tumor-stroma signaling circuit that links tumor inflammation to stromal remodeling and neutrophil infiltration in HNSCC.

Graphical abstract: **IL-1α drives tumor-stroma communication and neutrophil recruitment in HNSCC**. Here, we describe a mechanism in HNSCC that promotes high tumor-associated neutrophil (TAN) density, a biomarker associated with poor prognosis. Tumor-derived IL-1α activates stromal cells to adopt an inflammatory phenotype, resulting in the release of CXCL8, GM-CSF, and G-CSF to enhance neutrophil recruitment, activation, and survival. *IL1A*-positive tumor islets are surrounded by *CXCL8/CSF3*-rich stroma with elevated TAN densities in both tumor and stromal compartments. These TANs exhibit increased frequencies of MPO– and NE-positive cells, revealing a spatially organized inflammatory tumor microenvironment. Figure was created using BioRender.

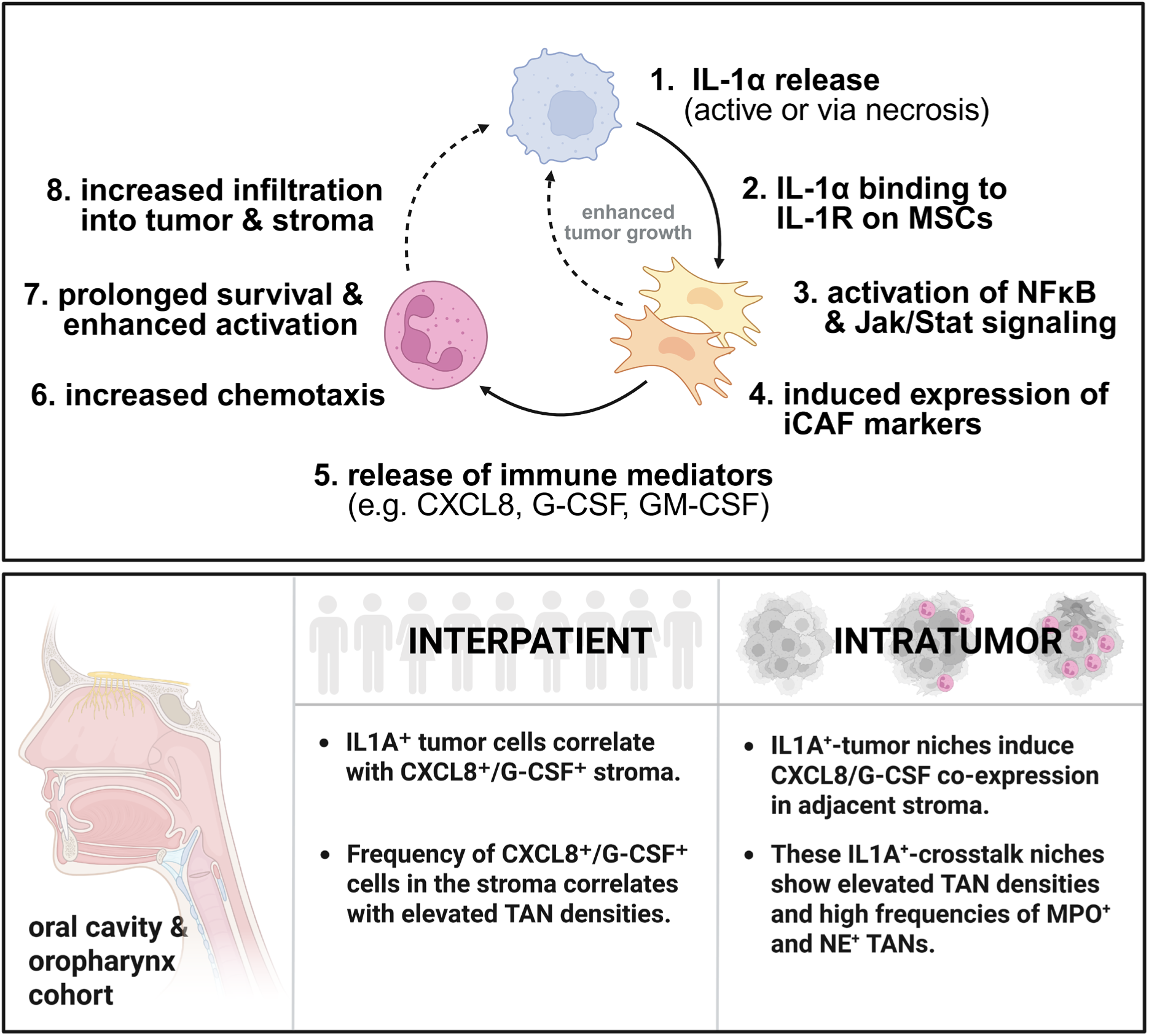

## Introduction

Head and neck squamous cell carcinoma (HNSCC) ranks among the most frequent and aggressive malignancies worldwide and is commonly diagnosed at advanced stages. Despite advances in surgery, radiotherapy, chemoradiation, and immunotherapy, the 5-year survival rate has only modestly improved^1^. A major reason lies in the biological heterogeneity of the tumor microenvironment (TME), which critically determines disease progression, immune evasion, and therapeutic response. In HNSCC, the TME consists of malignant epithelial cells, stromal fibroblasts, mesenchymal stromal cells (MSCs), endothelial cells, and diverse immune populations that interact via complex cytokine and chemokine networks^2,3^. These reciprocal interactions generate an inflammatory, angiogenic, and immunosuppressive milieu that fuels tumor growth. In particular, the paracrine crosstalk between cancer cells and fibroblastic stromal cells has emerged as a key driver of tumor progression and immune modulation^4^. Upon exposure to tumor-derived factors, MSCs can undergo transcriptional and epigenetic reprogramming into cancer-associated fibroblasts (CAFs)^5^, which remodel the extracellular matrix and release immunoregulatory mediators that shape leukocyte recruitment and function^6^. IL-1 has been identified as a key upstream signal driving inflammatory CAF differentiation^7–9^, and IL-1 driven stromal programs may influence immune cell recruitment, including neutrophils. At the tissue level, dynamic exchange between tumor and stromal compartments generates spatially and functionally heterogenous microenvironments that locally influence central tumor biological processes including tumor invasion and metastasis^10–13^. How these microenvironmental regions – or the intratumoral niches they form – affect tumor cell biology is relatively well understood; however, their impact on immune cell recruitment is less clear.

TANs have emerged as critical regulators of the TME. In HNSCC and other solid malignancies, neutrophils exhibit functional alterations in the blood and within tumors^14,15^. A high preoperative intratumoral TAN density has been identified as a robust predictor of poor outcome. While the clinical and prognostic relevance of TANs is firmly established, the precise mechanisms directing their recruitment to tumors and their subsequent intratumoral localization and functional adaptation remain incompletely understood. Classical recruitment pathways, including chemokine gradients mediated by CXCR1/2 ligands (e.g. CXCL1, CXCL2, CXCL8), are operative in inflammation and cancer and are complemented by cytokines and growth factors including G-CSF, GM-CSF, IL-6, type I interferons, TNFα and TGFβ (reviewed in^16^). Additional tumor– and stroma-derived signals, such as damage-associated molecular patterns (S100A8/S100A9, HMGB1), lipid mediators (such as PGE2, LTB4), and hypoxia-driven factors (HIF-1α, VEGF), further shape TAN trafficking and retention^17–19^. Adhesion molecules on neutrophils and endothelial cells, such as selectins, ICAM-1 and β2 integrins, facilitate rolling, extravasation and trans endothelial migration to the tumor site^20–22^. Spatial analyses in HNSCC indicate that the majority of TANs localizes to the tumor stroma rather than the tumor islets, positioning them to interact with stromal cells, vasculature, and extracellular matrix components, which likely influences their functional polarization^23,24^. Consistently, experimental imaging studies reveal that neutrophil recruitment and positioning within tumors are spatially regulated, with distinct stromal and intratumoral TAN populations exhibiting differential migratory behavior^24^. Upon recruitment, TANs can release granule-associated effectors such as myeloperoxidase (MPO) and neutrophil elastase (NE) through degranulation, contributing to extracellular matrix remodeling, modulation of signaling pathways, and generation of reactive species that further shape the TME^25,26^. Beyond their antimicrobial and effector functions, neutrophils within tumors exhibit remarkable plasticity. Single-cell and spatial transcriptomic studies reveal that TAN plasticity is not defined by distinct static states but by continuous transitions in activation and maturation^27,28^. Functionally, TANs can support anti-tumor immunity through cytotoxicity or NET release but are more often co-opted by tumors to promote angiogenesis, extracellular matrix (ECM) remodeling, immunosuppression and metastatic seeding^28–30^.

Here we investigated how spatially organized tumor-stroma crosstalk mediates the recruitment of TANs. Our study has been stimulated by emerging evidence indicating that IL-1-driven inflammatory signaling in CAFs enhances neutrophil-recruiting chemokines and growth factors, suggesting a potent stromal route for TAN modulation^31,32^. For instance, IL-1α signaling through IL-1 receptor type 1 (IL-1R1) on prostate MSCs induces a distinct inflammatory CAF (iCAF) phenotype *in vitro*, characterized by the expression of *IL-6, CXCL8, CSF3*, and other pro-inflammatory mediators^33^. In this study we demonstrate that IL-1α released from viable or necrotic tumor cells acts as a central upstream signal that reprograms patient-derived MSCs into an inflammatory CAF-like phenotype. These activated MSCs secrete neutrophil-recruiting and –survival factors such as CXCL8, G-CSF, and GM-CSF, establishing a tumor-stroma amplification loop that sustains TAN infiltration and activation. Using 3D co-culture systems, zebrafish xenografts, and spatially resolved analyses of HNSCC patient tissues, we show that *IL1A^+^* tumor islets are surrounded by *CXCL8/CSF3*-co-expressing cells of the stroma. These inflammatory niches are enriched in TANs across both tumor and stromal compartments.

This study defines a clinically relevant signaling circuit that integrates tumor inflammation, stromal remodeling, and TAN infiltration, highlighting IL-1α driven tumor-stroma crosstalk as a central orchestrator of neutrophil-rich inflammatory intratumoral niches.

## Results

### Establishment of systems that mimic tumor and stroma communication and their influence on neutrophil recruitment

To confirm the clinical relevance of TAN in our HNSCC cohort, we quantified TANs in primary tumors from 83 patients with oropharyngeal and oral cavity cancer (Supp. Table 1a). High overall TAN density was significantly associated with reduced overall survival (p = 0.008) (Fig. 1a), underscoring the prognostic impact of neutrophil infiltration in HNSCC. Notably, our cohort reflected the typical male predominance of HNSCC (2.7:1 ratio). The survival association reached statistical significance in males, while a similar trend was observed in females, suggesting that the lack of statistical relevance in the female subgroup may reflect limited statistical power rather than true sex-specific divergence in TAN biology (not shown). Spatial mapping of CD66b^+^ TANs across histologically defined tumor regions revealed a conserved pattern in both men and women: TANs were most abundant in the stroma, intermediate at the tumor-stroma interface, and lowest in the tumor core (Fig. 1b), indicating that TAN infiltration patterns are spatially structured and stromal-dominant. TAN distribution across compartments did not differ between sexes.

**Figure 1:**
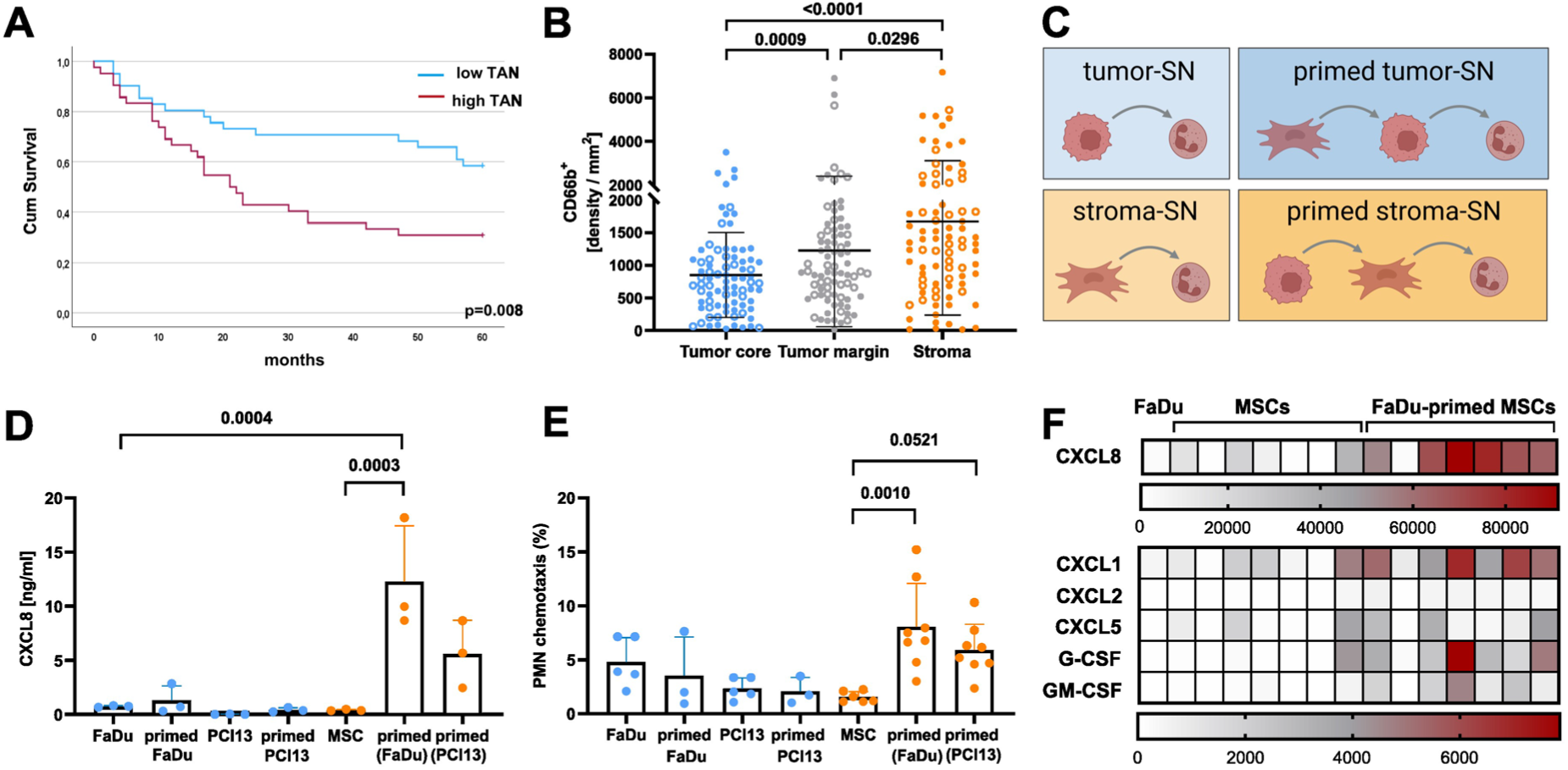
Tumor-stroma communication modulates neutrophil recruitment and density. (A) Kaplan-Meier analysis showing that neutrophil density negatively correlates with patient outcome, n=83 patients. (B) CD66b^+^ TAN were analyzed by immunofluorescence and quantified in tumor, tumor margin and stroma of HNSCC patients (mean ± SD, n=83, circles = males, open circles = females). (C) Schematic overview of the *in vitro* system for tumor-stroma communication, generated using BioRender. (D) SNs from tumor (FaDu, PCI13), MSC-primed tumor, MSCs and tumor-primed MSCs were analysed for released CXCL8 by ELISA, mean + SD, n=3. (E) The impact of these SN on PMN chemotaxis was analyzed using transwell assays, mean + SD, n=3-8. (F) Luminex screening was performed on SNs from FaDu cells and seven patient-derived MSC lines, analyzed both in their naïve state and after FaDu-priming. CXCL8 was measured using ELISA. Data are depicted in pg/mL. Statistical analysis was performed with Kruskal-Wallis (B) and ordinary one-way ANOVA (D, E). *P*-values are indicated.

To mechanistically dissect these compartment-specific differences in TAN densities and the role of tumor-stroma communication in neutrophil recruitment, we established a complementary *in vitro* system. We analyzed the supernatant (SN) of tumor and patient-derived MSCs, as well as stroma-primed tumor and tumor-primed MSCs (Fig. 1c). When MSCs were primed with conditioned medium of two HNSCC cell lines (FaDu, PCl13), they secreted markedly higher levels of CXCL8 compared to naïve MSCs or tumor cells. In contrast, tumor cells exposed to MSC-conditioned medium did not show a similar increase (Fig. 1d). To model neutrophil responses *in vitro*, primary polymorphonuclear neutrophils (PMNs, i.e., neutrophils) were exposed to SNs from tumor-primed MSCs, which induced a pronounced chemotactic response in transwell assays, linking stromal CXCL8 induction to neutrophil attraction (Fig. 1e). To determine whether additional neutrophil-regulating factors were co-induced, we performed a Luminex multiplex screening of SNs derived from FaDu cells, different patient-derived MSC lines, and their FaDu-primed counterparts. Tumor-conditioned MSCs secreted high levels of chemokines and growth factors, including CXCL1, CXCL2, CXCL5, G-CSF, and GM-CSF, with CXCL8 showing the highest overall concentrations (Fig. 1f).

Together, these results demonstrate that tumor-derived factors reprogram stromal MSCs to augment the release of neutrophil attracting chemokines and pro-survival growth factors, providing a first mechanistic basis for the stromal enrichment of TANs observed in HNSCC patient samples.

### Tumor-derived IL-1α drives tumor-stroma crosstalk

To identify which tumor-derived factors are responsible for MSC priming, we analyzed tumor-conditioned media for released factors known to regulate CXCL8 and/or G-CSF. To ensure that measured CXCL8 originated exclusively from stromal cells, we generated a stable CXCL8-knockout (CXCL8-KO) FaDu tumor cell line which was integrated in subsequent experiments. Luminex screening of the secretome of MSCs primed with FaDu or FaDu CXCL8-KO SNs revealed consistent detection of IL-1α, IFNγ, MIF, and VEGF across tumor SN batches. Their concentrations correlated with one another and with MSC-derived CXCL8 and/ or G-CSF (Fig. 2a, b, Supp. Fig. 1a, b). IL-1β and TNFα were not detected (Supp. Fig. 1b). As IFNγ, MIF, and VEGF are transcriptionally regulated downstream of IL-1α-NF-κB signaling, these data suggest IL-1α as a potential upstream regulator of tumor autocrine signaling and a key mediator of tumor-induced stromal priming. Next, we confirmed a dose-dependent induction of CXCL8 and G-CSF in MSCs by recombinant IL-1α (Fig. 2c). Conversely, neutralization of tumor-derived IL-1α during MSC stimulation with FaDu CXCL8KO-derived SN effectively suppressed the induction of both factors (Supp. Fig. 1c), confirming IL-1α as the key soluble mediator which induces CXCL8 and G-CSF in stromal cells.

**Figure 2:**
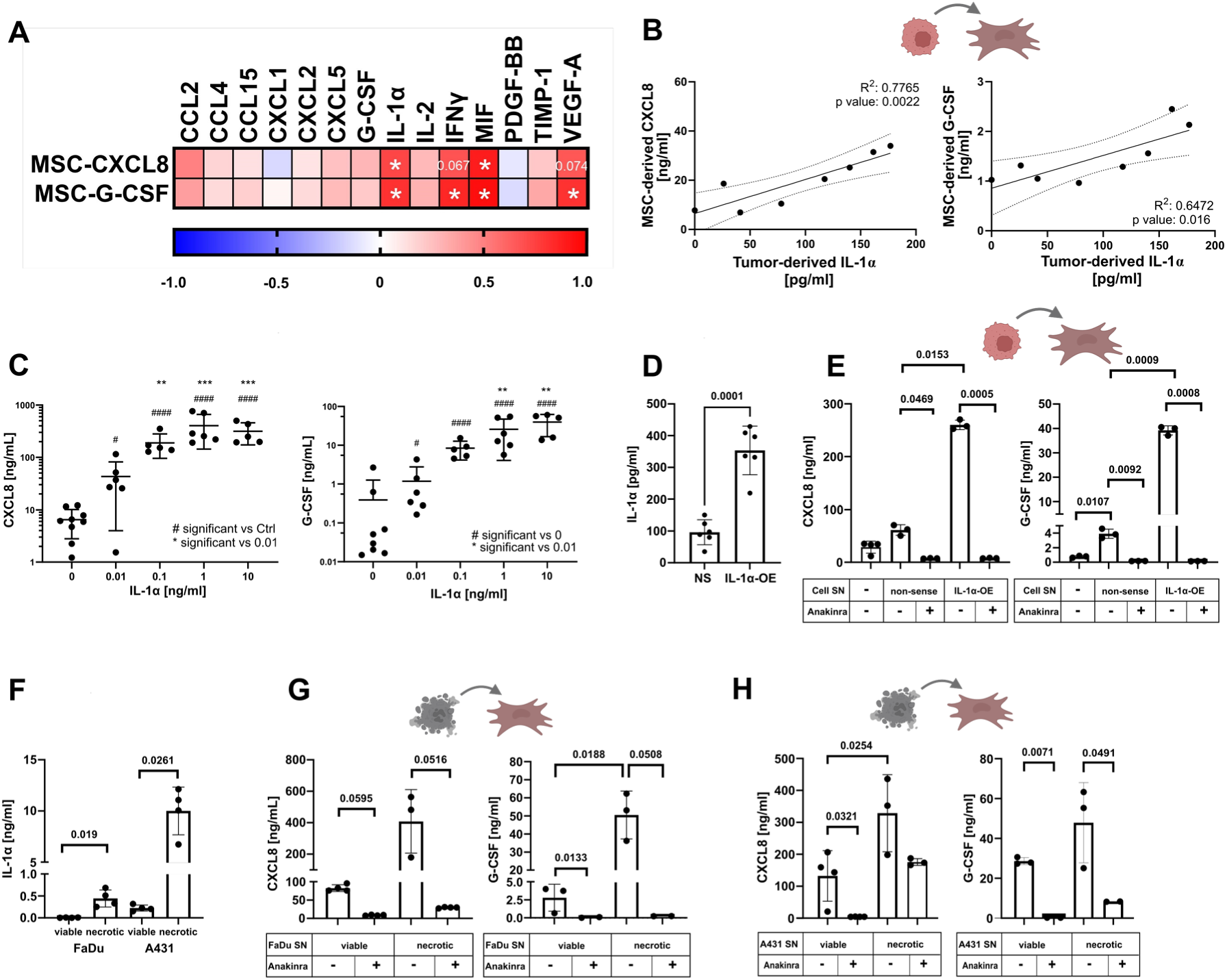
Tumor-derived IL-1α induces CXCL8 & G-CSF release by stroma cells. (A) MSCs were stimulated for 24 h with FaDu tumor-conditioned SNs. Tumor-derived cytokines (columns) were quantified by Luminex, and MSC-derived CXCL8 and G-CSF (rows) by ELISA. Data are displayed as a correlation matrix. (B) MSCs were treated with FaDu tumor SNs for 24 h. Tumor-derived factors were analyzed by Luminex; MSC-derived CXCL8 and G-CSF were quantified by ELISA. Tumor-derived IL-1α correlates with MSC-derived CXCL8 and G-CSF. (C) MSCs were treated with recombinant IL-1α for 24 h. Release of CXCL8 and G-CSF was analyzed by ELISA. (D) IL-1α was measured in control (non-sense, NS) and IL-1α overexpressing FaDu cells (IL-1α-OE) using Luminex. (E) MSCs were treated with SNs from non-sense and IL-1α-OE cells. MSC-derived CXCL8 and G-CSF were quantified by ELISA. (F) IL-1α release was determined in the SN of viable and necrotic FaDu and A431 cells. MSCs were treated with the SN of viable and necrotic FaDu (G) or A431 (H) cells for 24 h. MSC-derived CXCL8 and G-CSF were quantified by ELISA. Statistical significance was assessed after log-transformation using an ordinary one-way ANOVA with Tukey’s multiple comparisons test (C), while paired *t*-tests were applied for panels D-H. Data are shown as mean ± SD. In panel C, significance levels are indicated as # or * (p ≤ 0.05), ## or ** (p ≤ 0.01), ### or *** (p ≤ 0.001), and #### or **** (p ≤ 0.0001); all other p values are shown numerically. Symbols (BioRender) are included to show the origin of the analyzed SNs.

Since IL-1α can be actively secreted or passively released during necrotic cell death, we explored both sources. IL-1α-overexpressing FaDu (IL-1α-OE) cells, generated for this study, secreted moderately elevated IL-1α levels of up to 400 pg/mL compared to non-sense (NS) control cells (Fig. 2d). The increase in IL-1α was not associated with changes in overall IFNγ, MIF, or VEGF levels, nor did it alter chemokine or growth factor secretion by IL-1α-overexpressing FaDu cells (Supp. Fig. 1d,e). NS-FaDu SNs induced CXCL8 and G-CSF release from MSCs, which was efficiently blocked by the IL-1 receptor antagonist Anakinra, confirming tumor-derived IL-1α as the key mediator in this tumor-stroma communication. SNs from IL-1α-OE FaDu cells further enhanced CXCL8 and G-CSF release by MSCs (Fig. 2e). Moreover, SN from necrotic FaDu cells, containing moderate levels of IL-1α (Fig. 2f), strongly stimulated CXCL8 and G-CSF release from MSCs in an IL-1R-dependent manner (Fig. 2g), an effect that was also observed with A431-derived SN, supporting the general relevance of IL-1α-mediated tumor-stroma communication across carcinoma types (Fig. 2f,h).

Together, these data establish IL-1α, whether secreted by viable tumor cells or released upon necrosis, as a central driver of G-CSF and CXCL8 production via tumor-stroma communication, highlighting stromal paracrine signaling as a major source of neutrophil-regulating factors in the TME. To standardize subsequent stimulation conditions, the IL-1α concentration–response data (Fig. 2c) were used to define physiologically relevant and maximal activation ranges. FaDu non-sense and IL-1α-OE SNs released moderate IL-1α levels, comparable to those detected in our tumor-stroma models, whereas recombinant IL-1α at 10 ng/mL represents the plateau of stromal activation achievable during tumor cell necrosis.

### IL-1α induces an inflammatory CAF-phenotype in mesenchymal stromal cells

To uncover the cell biological program underlying the observed cytokine production, we performed bulk RNA sequencing of MSCs following IL-1α stimulation for 6 and 24 hours (Fig. 3, Supp. Fig. 2). IL-1α treatment triggered extensive transcriptional reprogramming (Fig. 3a, Supp. Fig. 2a,b), with 528 genes upregulated and 435 genes downregulated by ≥2-fold. Differentially expressed gene (DEG) clustering (adjusted p < 0.05) revealed marked temporal shifts. We examined the expression of selected myofibroblast CAF (myCAF), iCAF, and antigen-presenting CAF (apCAF) marker genes^34,35^. Canonical mesenchymal markers (*VIM, PDGFRA*) remained stably expressed, whereas myCAF-associated genes (*COL11A1, MMP11, MYL9*) were downregulated. *ACTA2* (encoding αSMA) was unchanged. In contrast, iCAF-related genes such as *PTGS2, IL6, CXCL1, CXCL2*, and *CSF3* were strongly induced, along with transcripts linked to inflammation, angiogenesis, and ECM remodeling (Fig. 3a, Supp. Fig. 2a,b). Matrisome analysis^36^ identified 75 upregulated matrisome-associated genes (≥ 3-fold), including 11 core matrisome components and 64 ECM regulators, affiliated proteins, or secreted factors, whereas 20 matrisome genes were strongly repressed by IL-1α-stimulation (≤ –3-fold) (Supp. Table 2). Hallmark pathway enrichment (Fig. 3b, Supp. Fig. 2c) and STRING protein network analysis (Fig. 3c) confirmed activation of ECM remodeling, epithelial-mesenchymal transition (EMT) and inflammatory signaling particularly IL-1, JAK/STAT, and NF-κB pathways. To assess whether the IL-1α-induced transcriptional program was conserved across mesenchymal stromal cells, we performed 6 h IL-1α stimulation and bulk RNA sequencing in a second independent MSC line. Comparison of significantly upregulated genes revealed a substantial overlap between both cell lines, indicating that the inflammatory CAF-like program is not restricted to a single MSC line (Supp. Fig. 2d).

**Figure 3:**
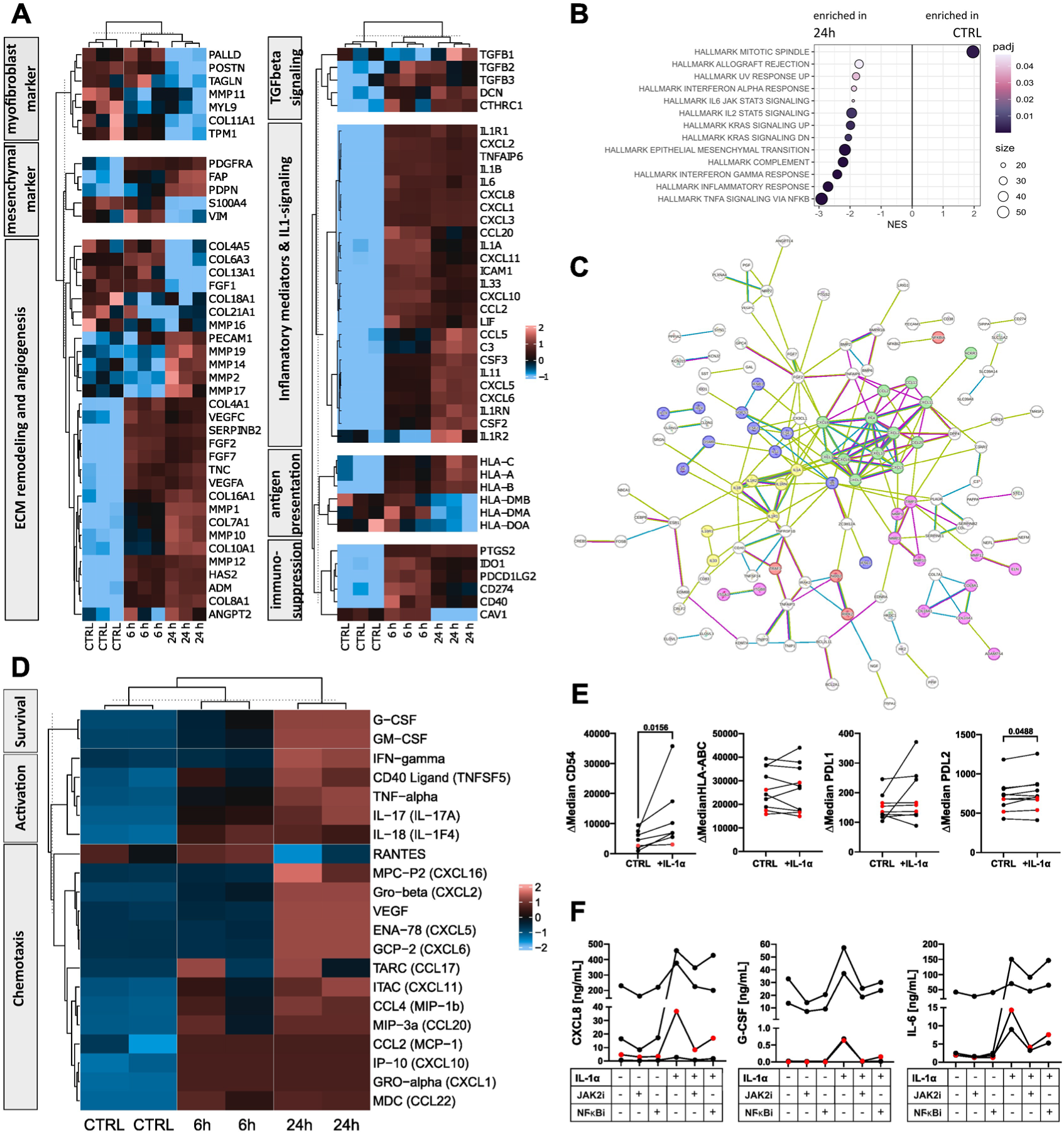
IL-1α treatment induces an inflammatory CAF-phenotype in MSCs. MSCs were treated with recombinant IL-1α for 6 and 24 h. Gene expression was analyzed by Bulk RNAseq. (A) Heatmap depicts functional DEGs linked to the biology and function of CAFs. (B) Enriched hallmark pathways are shown for untreated (ctrl) and 24 h IL-1α-treated MSCs. C) Pathway analysis (STRING v12.0) highlights local network cluster: IL-1 receptor activity and IL-1 family (yellow), chemokine receptors bind chemokines (green), extracellular matrix remodeling (pink), JAK-STAT (blue) and NFκB (red) signaling pathways. (D) Release of cytokines, chemokines and growth factors was analyzed by Luminex. Heatmap depicts proteins functioning in PMN survival, activation and chemotaxis. (E) Surface markers were analyzed by flow cytometry in different MSCs. (F) Inhibition of the Jak/Stat pathway by AZD1480 (JAK2i) or NFκB by Activation Inhibitor III (NFκBi) inhibit the release of CXCL8, G-CSF and IL6 in different MSCs. Red dots in E and F indicate the MSC cell line that was used for A-D. Statistical analysis was performed with paired t-tests. *P*-values are depicted.

Validation by RNAscope (*CXCL8, CSF3*) and COX-2 immunostaining confirmed data from bulk RNA sequencing (Supp. Fig. 2e). Luminex and ELISA analyses furthermore demonstrated significantly increased secretion of pro-inflammatory cytokines, growth factors and matrisome-associated proteins, including CXCL1, 2, 5 and 8, CCL20, IL-1β, IL-6, G-CSF, GM-CSF, and VEGF on the secretome level (Fig. 3d, Supp. Table 1b). Flow cytometry analysis revealed no induction of surface expression of HLA-A, –B, or –C despite transcriptional upregulation, and HLA-DR remained undetectable (data not shown), excluding an apCAF-like differentiation. Instead, CD54 (ICAM-1) and PD-L2 were consistently upregulated, while PD-L1 levels were unchanged across MSC lines (Fig. 3e). Finally, pharmacological inhibition of JAK/STAT and NF-κB signaling abrogated IL-1α-induced release of CXCL8 and G-CSF (Fig. 3f), confirming these pathways as essential mediators of the IL-1α-driven iCAF-like transcriptional program.

It has been shown that inflammatory CAFs can modulate and support tumor growth^37^. To investigate whether IL-1α-stimulated MSCs acquire a functionally relevant iCAF-like program, we next assessed the impact of their conditioned medium on FaDu tumor spheroids. Spheroid growth was significantly enhanced by SN from IL-1α-stimulated MSCs (Supp. Fig. 2f), providing functional evidence that IL-1α-driven iCAF-like MSCs can promote tumor growth. Together, these data demonstrate that IL-1α induces an inflammatory CAF-like transcriptional and secretory program in resting MSCs, characterized by elevated cytokines, growth factors, and ECM regulators known to promote tumor progression. We therefore refer to those stimulated stromal cells as IL-1α-iCAF-like MSCs.

### IL-1α-induced iCAF-like signature correlates with distinct CAF subtypes in HNSCC and poor prognosis

To assess the disease-related relevance of the IL-1α-induced iCAF-like program, we generated a gene signature based on all significantly upregulated genes in IL-1α-treated MSCs (Supp. Table 2). Gene set enrichment analysis (GSEA) revealed significant overlap with previously described HNSCC CAF subtypes. Zhang et al. identified eight CAF subclusters in HNSCC^38^. Our IL-1α-iCAF-like signature was enriched in clusters 3, 4, and 6, which are linked to ECM remodeling, EMT-related genes, and antigen presentation, and were associated with poorer overall survival (Fig. 4a). Obradovic et al. defined five molecularly distinct CAF clusters (HNCAF-0 to HNCAF-4)^39^. Our IL-1α-iCAF-like signature was enriched in HNCAF-1 (iCAF), HNCAF-3 (iCAF-like, ECM remodeling), and HNCAF-0 (iCAF), but not in HNCAF-2 (myCAF), which was, in fact, enriched in untreated control MSCs (Fig. 4b). Notably, HNCAF-1 has been linked to immunosuppression and poor overall survival, whereas HNCAF-0 and HNCAF-3 have been linked to favourable responses to anti-PD-1 immunotherapy. Choi et al. reported five CAF clusters (CF0–CF4) including an iCAF CF1/CXCL8 cluster representing the most progressed fibroblast state^40^. Our IL-1α-treated MSCs upregulated all five CF1 marker genes (CXCL8, IL24, AKR1B1, MMP3, MMP1) 2–8 fold, whereas markers of other clusters were unchanged or downregulated (except CTHRC1 from CF0, 2-fold upregulated) (Figure 4c, Supp. Table 2).

**Figure 4:**
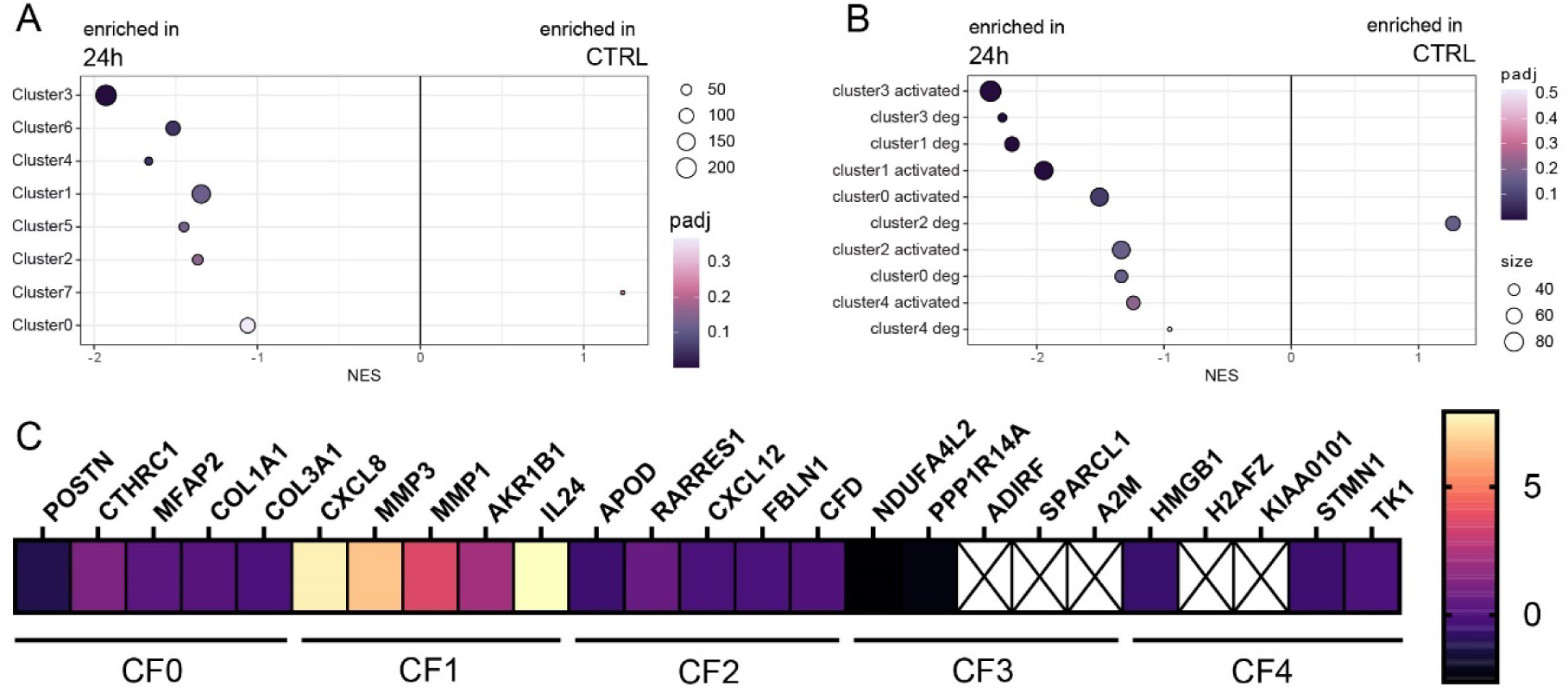
IL-1α-induced MSC signature aligns with described CAF clusters in HNSCC. GSEA was performed on genes differentially expressed in MSCs after 24 h IL-1α treatment versus untreated controls. (A) Enrichment analysis of the IL-1α signature across eight CAF clusters^38^. Clusters 3, 4, and 6 are associated with ECM remodeling, EMT, antigen presentation, and worse overall survival. (B) Enrichment analysis across five CAF clusters^39^. HNCAF-0 (iCAF), HNCAF-3 (iCAF-like/ECM remodeling), and HNCAF-1 (iCAF, immunosuppressive) show high similarity to the IL-1α program; HNCAF-1 is linked to poor prognosis, whereas HNCAF-0/3 have been associated with favorable responses to anti-PD-1 immunotherapy. (C) The heatmap shows log2 fold changes of published CAF cluster marker genes^40^ in IL-1α-treated MSCs relative to control. The CF1/CXCL8 cluster represents an iCAF cluster and the most progressed fibroblast state.

Collectively, these analyses link the IL-1α-induced iCAF-like program in MSCs to clinically relevant CAF subtypes associated with ECM remodeling, immunosuppression, and poor patient prognosis. Importantly, mapping to HNCAF-0 and HNCAF-3 suggests that this program may also influence responses to immune checkpoint blockade, further supporting the translational relevance of our *in vitro* findings.

### MSC-derived factors modulate PMN survival and activation

Building on our observation that IL-1α-primed MSCs adopt an inflammatory CAF-like program along with neutrophil recruitment and survival factors, we next investigated how stromal signals actually affect neutrophil survival and activation. Primary human PMNs were exposed to control SNs from non-sense (NS) or IL-1α-overexpressing (OE) FaDu cells, or to SNs from MSCs preconditioned with such tumor-derived factors. Recombinant IL-1α was used to assess direct effects. SNs from IL-1α-OE tumor cells alone did not affect PMN survival compared to NS controls (Fig. 5a). In contrast, SNs from tumor-primed stromal cells significantly enhanced PMN survival, with the strongest effect observed after priming with IL-1α-OE FaDu SN (Fig. 5a). Recombinant IL-1α alone had no effect (not shown), indicating that MSC-mediated secondary signals are required. Neutralization experiments suggested that GM-CSF, rather than G-CSF contributed to enhanced PMN survival (Fig. 5b).

**Figure 5:**
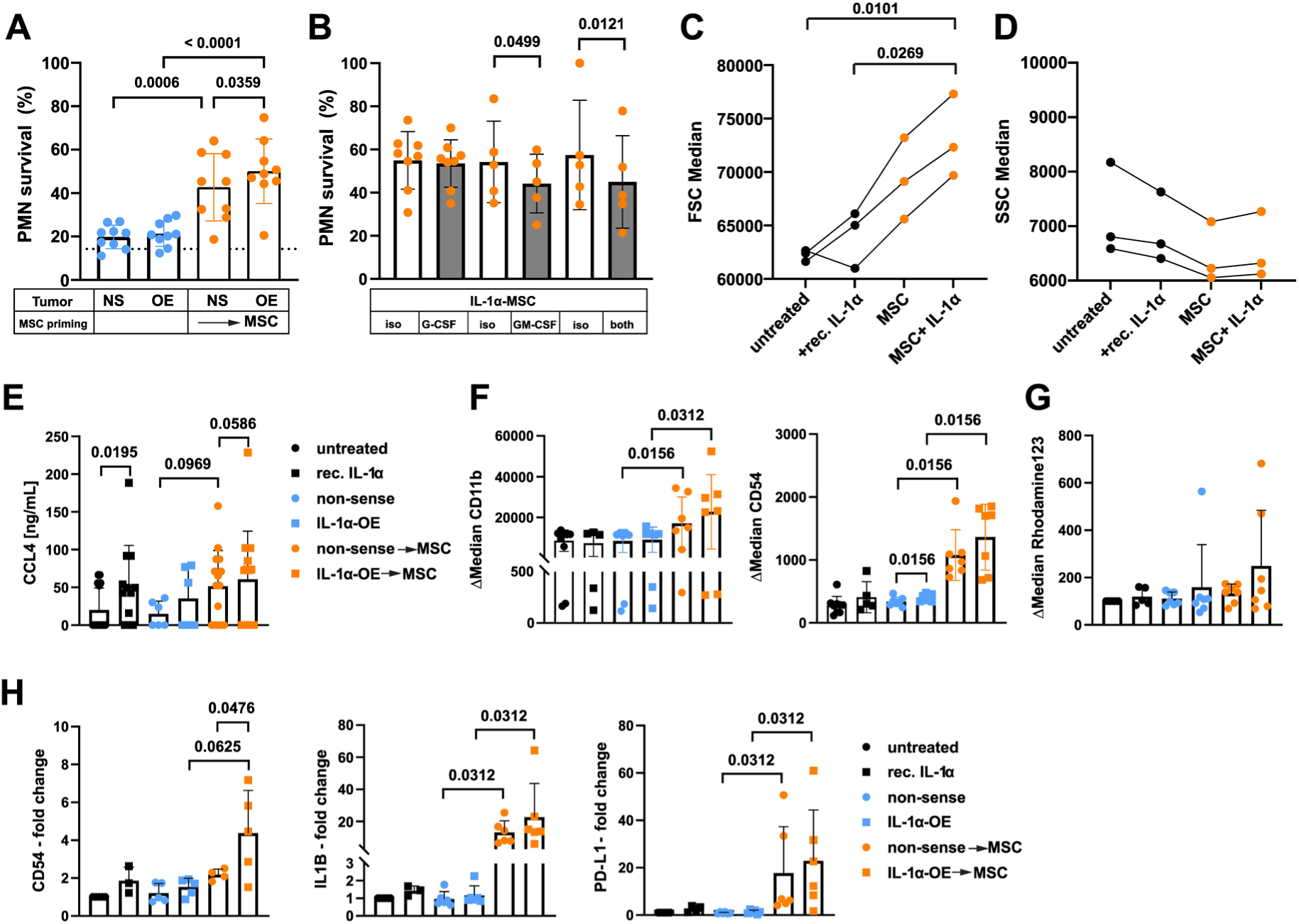
MSC-derived factors modulate PMN survival and activation. PMNs were treated with recombinant IL-1α and SNs from non-sense (NS) or IL-1α-overexpressing (OE) FaDu cells, as well as from MSCs preconditioned with these tumor cell SNs. A) PMN survival was analyzed by Annexin V/7-AAD staining (dotted line indicates the mean of untreated PMNs and (B) in the presence of neutralizing antibodies against G-CSF, GM-CSF, or both. (C,D) Activation was assessed using flow cytometry, determining changes in cell size (C) and granularity (D). (E) CCL4 release was quantified by ELISA. (F) Surface activation marker CD11b and CD54 were analyzed by flow cytometry. (G) ROS productionwas measured using 123-DiRhodamine. (H) Expression of marker genes was assessed by qRT-PCR. Statistical analysis was performed with paired t-tests (A,B), ordinary one-way ANOVA with Tukey’s multiple comparisons test (C) or Wilcoxon signed rank test (E-H). Data are plotted as mean + SD (E, G, H) or mean ± SD (A,B,F). *P*-values are stated.

We next explored effects of IL-1α and primed stromal cells on PMN activation. PMNs exposed to MSC SNs showed increased cell size without changes in granularity (Fig. 5c,d), enhanced CCL4 release (Fig. 5e), and upregulation of CD11b and CD54 (Fig. 5f). At the transcriptional level, tumor-stroma communication strongly induced *IL1B*, *PD-L1*, and *CD54*, with *CD54* most prominently increased in PMNs exposed to IL-1α-OE-primed MSC SN (Fig. 5h). SN from MSCs treated with recombinant IL-1α recapitulated key aspects of this activation phenotype, including increased CCL4 release and trends toward elevated CD11b and CD54 surface protein levels, as well as induction of *IL1B, PD-L1* and *CD54* transcripts, although these changes did not consistently reach statistical significance (Supp. Fig. 3a,b,d). MSC-derived factors did not induce classical neutrophil effector functions, as neither FaDu-primed nor IL-1α-OE FaDu MSC SN triggered ROS production, degranulation, or release of MPO, MMP9, or arginase-1 (Fig. 5g and Supp. Fig. 3d–f). Despite inducing neutrophil activation in terms of cytokine responses and surface marker expression, IL-1α-treated MSCs failed to elicit degranulation-associated effector mechanisms, indicating that tumor-MSC crosstalk promotes a transcriptionally activated neutrophil state amplified by IL-1α-driven stromal signaling.

### MSC-derived factors enhance PMN chemotaxis and infiltration *in vitro* and *in vivo*

Having established that IL-1α-programmed MSCs promote PMN survival and activation, we next examined their role in recruitment. Consistent with cytokine and chemokine secretion profiles, primed stromal cells induced stronger PMN recruitment than tumor cells, an effect further enhanced by IL-1α overexpressing tumor cells (Fig. 6a). Neutralization of CXCL8 significantly reduced IL-1α-OE-MSC-induced PMN migration (Fig. 6b), whereas blocking other CXCR2 ligands had no detectable effect (data not shown). Accordingly, genetic deletion of CXCL8 in MSCs completely abolished IL-1α-induced chemotaxis, despite continued release of CXCL1, CXCL2, and CXCL5 (Fig. 6c,d). Together, these data identify CXCL8 as the dominant and non-redundant stromal chemokine mediating PMN recruitment.

**Figure 6:**
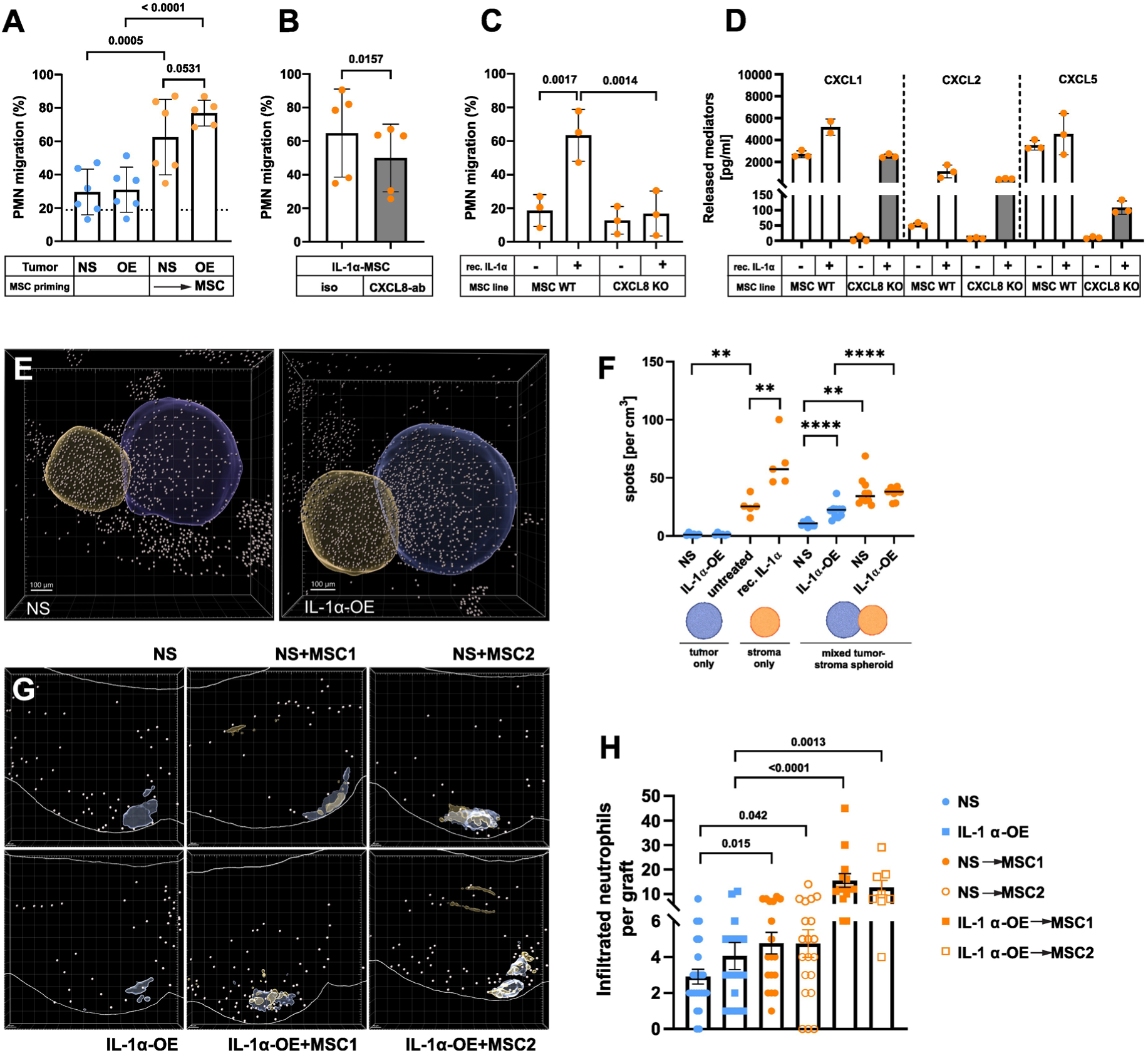
MSC-derived factors enhance PMN chemotaxis and infiltration, in vitro and in vivo. PMNs were treated with SN from non-sense (NS) or IL-1α-overexpressing (OE) FaDu cells, as well as from MSCs preconditioned with these tumor cell SN. (A) PMN migration toward tumor– and MSC-derived factors was analyzed by trans-well assays (dotted line indicates the mean of untreated PMNs) and (B) in the presence of neutralizing antibodies against CXCL8. (C) PMN migration was assessed by transwell assays toward SN collected from IL-1α–treated or untreated wild-type (WT) and CXCL8-knockout (KO) MSCs. (D) These SN were analyzed for released chemoattractants using Luminex. (E) Representative images of PMN recruitment into mixed NS FaDu-MSC or IL-1α OE-FaDu-MSC spheroids (tumor spheroids, blue; MSC spheroids, orange; PMNs, gray). (F) Quantification of PMN recruitment into tumor spheroids (NS or IL-1α-OE FaDu only), MSC spheroids (untreated vs. IL-1α-treated), and mixed tumor-MSC spheroids was analyzed by confocal microscopy (n = 6–10). (G) Representative images of FaDu cells (blue), MSCs (orange) and neutrophils (gray) 24 h post-injection into the perivitelline space of zebrafish larvae. Upper row: FaDu NS control cells; lower row: IL-1α-OE FaDu cells. Left panel: FaDu cells only; middle panel: FaDu cells + MSC1; right panel: FaDu cells + MSC2. (H) Quantification of infiltrated neutrophils per graft at 24 h post-injection. Statistical analysis was performed using paired *t*-tests (A-C), Mann–Whitney tests (F), or Wilcoxon signed-rank tests (H). Data are presented as mean ± SD (A-C), median (F), and mean ± SEM from three independent experiments (H). *P* values are indicated.

In the next series of experiments, we sought to extend the 2D cell culture experiments to 3D and zebrafish models. In 3D models, IL-1α-stimulated MSC spheroids showed robust PMN infiltration, whereas IL-1α-stimulated tumor-only spheroids did not. In mixed tumor-MSC spheroids, IL-1α overexpression increased PMN recruitment into both compartments (Fig. 6e,f).

In a zebrafish larvae xenograft model using Tg(lyz:dsred) embryos with fluorescent neutrophils (Fig. 6g,h), injection of NS or IL-1α-OE FaDu cells alone resulted in comparably low neutrophil infiltration. Co-injection of NS cells with MSC moderately increased neutrophil numbers. Co-injection of IL-1α-OE FaDu cells with MSC led to a pronounced increase in neutrophil infiltration. These findings demonstrate that tumor-derived IL-1α also markedly amplifies stromal cell-mediated neutrophil recruitment *in vivo*, reinforcing the role of IL-1α-induced stromal activation in driving TAN chemotaxis.

Together, these data demonstrate that IL-1α-activated stromal cells coordinate neutrophil survival, activation and recruitment, across *in vitro* and *in vivo* tumor models.

### IL-1α creates an inflamed and TAN enriched HNSCC tumor microenvironment

To assess the impact of IL-1α on the TME and TAN in patients, we performed spatial analyses on primary HNSCC tissues from 21 patients with oropharyngeal and oral cavity cancer (Supp. Table 1b). To this end we selected *IL1A, CXCL8*, and *CSF3* as representative markers of the IL-1α-induced stromal niche, reflecting the upstream cytokine (*IL1A*) and its downstream mediators (*CXCL8* and *CSF3*). Figure 7a shows a representative tumor to illustrate the tumor-stroma segmentation, probe signals, density heat map and zonal analysis capturing tumor-stroma communication that was used to define IL1A^−^ and IL1A^+^ niches (Fig. 7a, 8a). Tumor heterogeneity is visualized by boxplots depicting *IL1A*, *CXCL8*, *CSF3*, and *CXCL8-CSF3* double-positive cells in tumor and stromal compartments. Despite substantial interpatient heterogeneity relative frequencies of *IL1A*^+^ cells were significantly higher in the tumor region as compared to the stroma (Supp. Fig 3g). Across the patient cohort tumor *IL1A* expression correlated positively with the frequency of stromal *CXCL8/CSF3* double-positive cells (R² = 0.2171, p = 0.0027) (Fig. 7b). Sex stratified analysis suggested a stronger association in male patients (n=16, R² = 0.6377, p = 0.0029) (Fig. 7c), whereas no significant correlation was detected in female patients, who also exhibited a trend towards lower *IL1A* expression (n = 5, p = 0.0865, (Supp. Fig. 3h)). Sex stratified analyses were limited by the small number of female patients. Patient age showed no correlation with cytokine expression in the overall cohort or within sex groups (data not shown).

**Figure 7:**
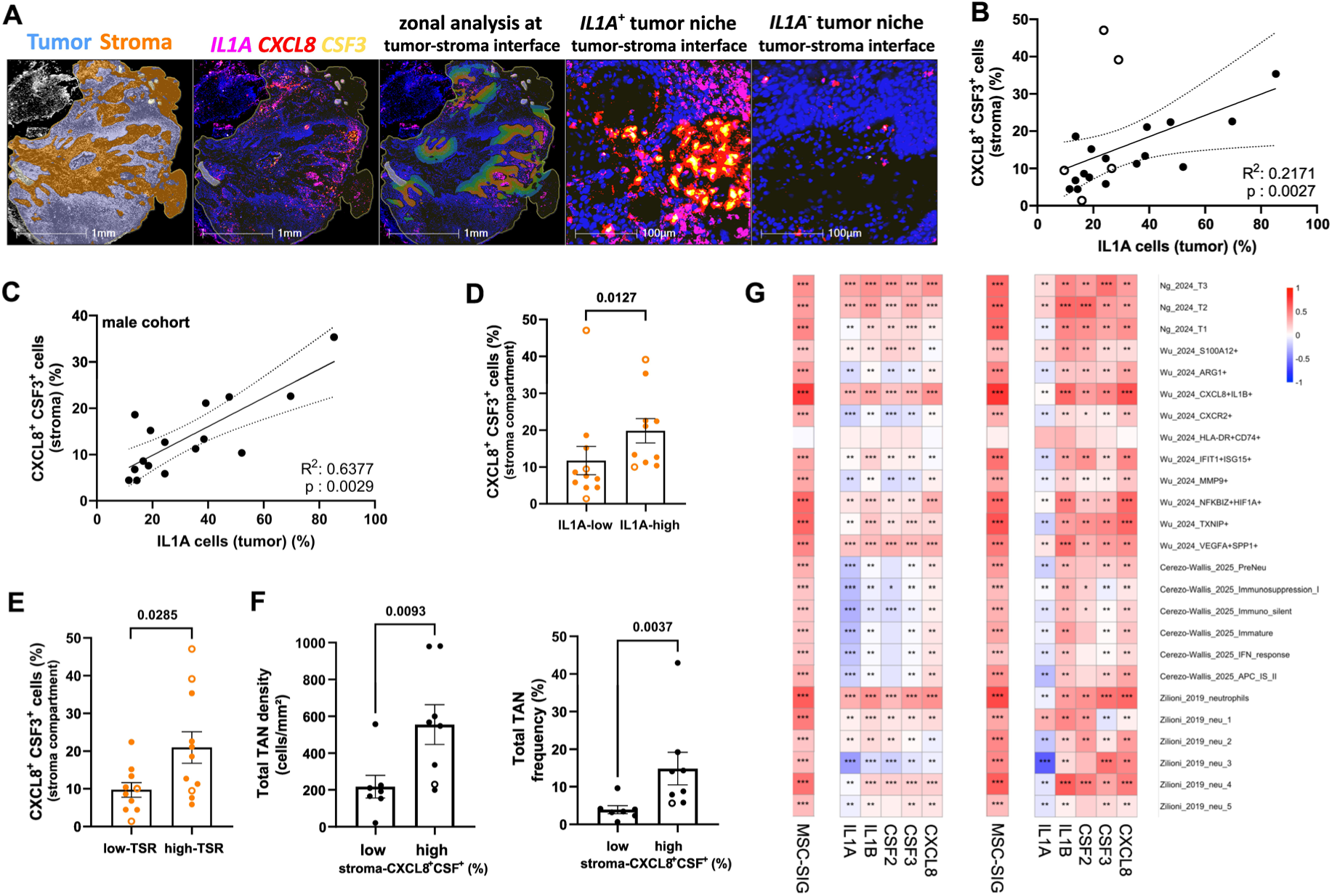
Interpatient analyses of IL1A, CXCL8, and CSF3 expression in primary HNSCC tumors and their relationship with TANs. (A) Representative tumor section illustrating tumor-stroma segmentation, RNAscope detection of *IL1A* (pink), *CXCL8* (red), and *CSF3* (yellow), and zonal analysis of tumor-stroma communication. Shown is also an example of an IL1A^+^ niche and an IL1A^−^ niche with corresponding RNA probe signals. (B,C) Correlation of tumor *IL1A* expression with stromal *CXCL8*^+^*CSF3*^+^ double-positive cells across the total cohort (B; n = 21; black circles = males, open circles = females) and in male patients only (C; n = 16). (D,E) Frequency of *CXCL8*^+^*CSF3^+^*double-positive stromal cells stratified by *IL1A*-negative vs. IL1A-positive tumors (D) and by TSR-low vs. TSR-high tumors (E). (F) Total TAN density and frequency in patients with low vs. high stromal *CXCL8*^+^*CSF3*^+^ cells. (G) Spearman correlation analysis of an IL-1α-induced MSC/iCAF gene signature (SIG) with published TAN signatures across TCGA HNSCC tumor samples (left, n = 515) and matched tumor-adjacent normal tissue (right, n = 44). The MSC/iCAF signature (MSC-SIG) was defined by the top 100 genes upregulated in MSCs following 24 h IL-1α stimulation. TAN signatures from four independent studies^41–44^ represent immature, inflammatory, and activated neutrophil populations. Individual cytokine and chemokine expression (*IL1A, IL1B, CSF2, CSF3, CXCL8*) shows concordant associations with multiple TAN subset signatures. Statistical analysis was performed using Spearman correlation (B,C,G) and Mann–Whitney test (D-F). Data are presented scatter plots (B,C), or mean ± SEM (E–J). *P* values are indicated (A-D). In (G) p values are indicated as follow: * p value < 0.05, ** p value < 0.01, *** p value <0.001.

To specifically uncover the role of *IL1A* in the TME, patients were divided by median tumor *IL1A* expression into *IL1A*-low and *IL1A*-high groups. Patients with *IL1A*-high tumors exhibited a significantly higher frequency of *CXCL8/CSF3* double-positive stromal cells (Fig. 7d). Stratification by tumor-stroma ratio (TSR ≤ 1.5 vs. > 1.5; (Fig. 7e)) revealed a comparable pattern, demonstrating that a higher TSR is associated with increased *CXCL8/CSF3* double positive cells in the stroma. When grouped by median stromal *CXCL8/CSF3* double-positive frequency, both, total TAN density and frequency (Fig. 7f) were significantly higher in the *CXCL8/CSF3*-high group. In contrast, high *IL1A* expression alone was not associated with increased abundance of TAN (data not shown).

These tissue-based analyses link tumor *IL1A* expression and stromal *CXCL8/CSF3* production to increased TAN abundance at the patient level. To determine whether this IL-1α–driven stromal–neutrophil axis is also evident in independent patient cohorts, we next interrogated public cancer patient transcriptomic datasets. We generated an IL-1α-induced MSC/iCAF like gene signature based on the top 100 genes upregulated in MSCs after IL-1α stimulation (Supp. Table 2).

We also extracted TAN subset signatures from four independent published studies^41–44^, spanning various transcriptional states. To assess whether the IL-1α-induced iCAF like transcriptional programs co-vary with TAN subset programs, we performed signature correlation analyses using HNSCC TCGA bulk RNA-seq data obtained from cBioPortal platform (https://www.cbioportal.org/^45–48^) without further deconvolution. With this approach, we observed robust positive correlations between the iCAF-like signature and nearly all TAN subset signatures with the exception of the HLA-DR^−^CD74^+^ subset described by Wu et al. (Fig 7g, left panel). Analysis of TCGA HNSCC tumor and adjacent tissue datasets revealed comparable or even stronger correlations in tumor-adjacent tissue, suggesting that this stromal-neutrophil association is not restricted to the TME (Fig. 7g). Notably, when analyzing individual cytokines, only *CXCL8* showed strong associations across TAN transcriptional states, whereas high *IL1A* transcript levels did not broadly correlate with increased TAN signatures. Despite this, *IL1A* transcript levels in HNSCC tumor tissue showed a stronger correlation with T2 and T3 TAN from Ng *et al.*, 2024^41^, CXCL8^+^IL1B^+^ and VEGFA^+^SPP1^+^ TAN from Wu *et al.*, 2024^42^ and neutrophil signatures from Zillionis *et al.*, 2019^44^, than in adjacent tissue. While these data are in line with a clear interconnection of stromal inflammation and TAN, our findings also prompted us to consider that *IL1A* and *IL1A*-induced stromal inflammation may act locally within discrete intratumoral niches rather than across the entire tumor; a possibility we next investigated through spatial mapping.

### IL-1α shapes a distinct stromal cytokine niche associated with TAN accumulation

To this end, we performed intratumoral spatial mapping in a subset of seven tumors. Tumor niches were defined based on *IL1A* expression, distinguishing *IL1A*^+^ and *IL1A*^−^ tumor islets along with their immediately adjacent stromal regions (200 µm) within the same tumor sample (Fig. 8a). *IL1A*^+^ tumor islets were consistently associated with increased frequencies of *CXCL8*^+^/*CSF3*^+^ double-positive cells in the surrounding stroma, indicating that the effect of *IL1A* on neutrophil regulating cytokines and growth factors is spatially confined (Fig. 8b). A zonal quantification within four concentric 50 µm-wide stromal zones away from the tumor-stroma border showed that *CXCL8/CSF3* double-positive cells were most enriched directly adjacent to *IL1A*^+^ tumor islets, underscoring the spatial coupling between tumor-derived IL-1α and stromal cytokine activation (Fig. 8c). We then assessed whether intratumoral niches with high cytokine expression exhibit increased TAN density and infiltration. Regions containing both tumor and stromal compartments were first stratified by *IL1A* expression in tumor islets using RNAscope signals, defining *IL1A*^−^ and *IL1A*^+^ niches. The same regions were then independently analyzed for TAN markers by cyclic immunofluorescence (MACSima) on consecutive slides (Fig. 8d). *IL1A*^+^ niches showed significantly higher TAN density than *IL1A^−^*regions in both tumor and stromal compartments (Fig. 8e,f). In contrast, CD3^+^ T-cell and CD68^+^ macrophage densities were not significantly affected by *IL1A* status (Fig. 8g,h), indicating a selective association between IL-1α-expressing tumor niches and neutrophil accumulation.

**Figure 8:**
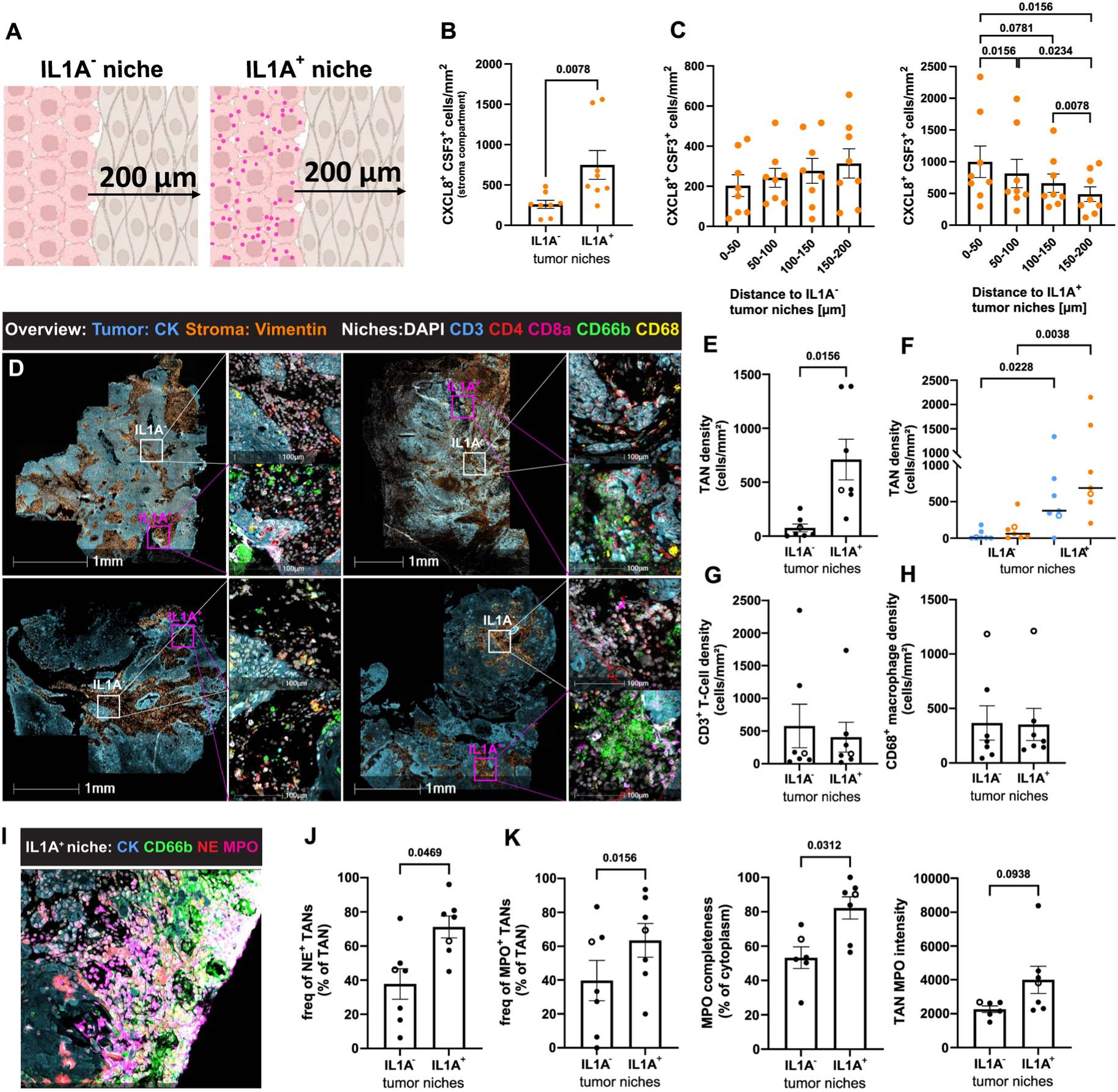
IL-1α-driven stromal inflammation is linked to neutrophil accumulation, spatial organization, and effector activation. (A,B) Densities of *CXCL8*^+^*CSF3*^+^ double-positive cells were quantified within 200 µm stromal regions adjacent to *IL1A*^−^ versus *IL1A^+^* tumor niches. Panel A shows the analysis scheme (created with BioRender). (C) Quantification of *CXCL8*^+^*CSF3*^+^ cell densities across four consecutive 50 µm stromal zones extending from *IL1A*^−^ or *IL1A*^+^ tumor islets (n = 8). (D) Representative images from four tumors stained for CK, VIM, CD66b, CD3, CD4, CD8, and CD68, including magnified views of *IL1A*^−^ (white) and *IL1A*^+^ tumor niches (pink), based on RNAscope analysis on consecutive slides (hereafter referred to as *IL1A* status). (E) Total TAN density within *IL1A*^−^ and *IL1A*^+^ tumor niches. (F) Spatial distribution of TANs across tumor (blue) and stromal (orange) compartments within these niches, stratified by *IL1A* status. (G) CD3^+^ T-cell density and (H) CD68^+^ macrophage density, stratified by *IL1A* status. (I) Representative image of an *IL1A*^+^ niche stained for CK, CD66b, NE, and MPO. (J,K) Frequency of NE^+^ TANs, and frequency of MPO^+^ TANs, cytoplasmic completeness, and total MPO intensity, stratified by *IL1A* status. Statistical analysis was performed using the Wilcoxon signed-rank test (B,C,E-H, J,K). Data are shown as mean ± SEM. *P* values are indicated (n = 7-8). CK, cytokeratin; VIM, vimentin; NE, neutrophil elastase; MPO, myeloperoxidase.

We next examined whether *IL1A*^+^ niches were associated not only with increased neutrophil presence but also with altered neutrophil effector characteristics. To this end we quantified the positivity of TANs for NE and MPO, two central effector molecules associated with neutrophil degranulation and NET formation. TANs within IL1A^+^ regions displayed a significantly higher frequency of NE positivity compared to *IL1A*^−^ niches (Fig. 8i,j). Similarly, MPO was elevated in *IL1A*^+^ regions, reflected by higher frequencies of MPO-high cells with an increased cytoplasmic completeness and total MPO intensity, indicating preservation of granule content and potential functional activity (Fig. 8i,k). In contrast, *IL1A*^−^ regions showed reduced MPO completeness, consistent with accumulation of terminally degranulated or NETotic TANs. Together, these data indicate that in patients with HNSCC, intratumoral *IL1A* locally instructs stromal inflammation, with consequences for neutrophil recruitment, accumulation, and phenotype in spatially confined niches. Our findings position IL-1α as a key organizer of a spatially confined stromal cytokine circuit that shapes both the abundance and local functional state of human TANs within cytokine-inflamed niches.

## Discussion

This study identifies a tumor-driven, stroma-amplified signaling circuit that governs neutrophil recruitment, spatial organization, and activation in HNSCC. By linking IL-1α-mediated tumor-stroma communication with TAN density and localization, we define how cytokine-driven stromal activation shapes the inflammatory landscape of the TME. IL-1α, released from viable or necrotic tumor cells, reprograms MSCs into inflammatory-like CAFs. These CAF secrete neutrophil-recruiting and survival factors, including CXCL8, G-CSF, and GM-CSF, forming a feed-forward loop that amplifies neutrophil infiltration and activation, with CXCL8 emerging as the dominant chemokine mediating TAN recruitment. This finding is clinically relevant, as elevated *CXCL8* expression has been linked to poor prognosis, increased recurrence, and aggressive disease in HNSCC patients^49,50^, underscoring its potential as both a prognostic biomarker and therapeutic target. GM-CSF on the other hand contributes to sustaining neutrophil viability and potentially proinflammatory polarization, consistent with its described role in extending neutrophil lifespan, inducing a proinflammatory transcriptional program and promoting NET formation^51,52^.

Tumor– and stroma-derived IL-1α has been implicated in the progression of multiple human cancers by promoting tumor growth and inducing vasculogenic factors, and by fostering tumorigenesis through NF-κB–dependent, senescence-associated stromal remodeling^53,54^. IL-1/IL-1R-dependent CAF populations have been observed across cancers, where they associate with immune modulation and therapy response^55^. IL-1α-induced iCAF-like MSCs are biased toward inflammatory and immune-modulating programs, and our matrisome analysis shows that they might also contribute to extracellular matrix production/ remodeling, distinguishing them from predominantly TGF-β-driven myCAFs, which primarily remodel the matrix and tissue organization^8^. CAF phenotypes are shaped by inflammatory versus fibrotic cues and remain plastic, adapting to microenvironmental changes associated with tumor evolution or treatment^37^. While IL-1α was the predominant initiating signal in our study, IL-1β can activate similar downstream pathways under defined conditions. Although recombinant IL-1β induced CXCL8 release in MSCs *in vitro* (data not shown), in our study *IL1B* transcripts were low and lacked the spatial patterning observed for *IL1A* in HNSCC tissue (data not shown), supporting IL-1α as the dominant spatial cue for localized stromal activation. IL-1α and IL-1β differ in their cellular release patterns; whereas IL-1β requires inflammasome-dependent proteolytic maturation to become active, IL-1α can be actively released, induced by hypoxia, and also functions as an alarmin when released from necrotic tumor cells^56,57^. Consistently, *IL1B* was induced in IL-1α-stimulated MSCs and in PMNs exposed to MSC-derived factors, highlighting dynamic tumor-stroma-neutrophil crosstalk. Previous studies have shown that IL-1α and IL-1β can initiate and reinforce inflammatory tumor niches, respectively^58,59^. Together, these findings suggest complementary but context-dependent roles for IL-1 family members in establishing and reinforcing inflammatory tumor niches, with IL-1α initiating spatially confined niches and IL-1β reinforcing inflammatory signaling. Different from reports describing direct IL-1 effects on neutrophils in sterile inflammation, our data indicate that IL-1α primarily acts indirectly via stromal cells in the tumor setting. Direct IL-1α exposure did not affect PMN survival or migration, consistent with previous reports for IL-1^60–62^, whereas MSC-derived factors robustly promoted neutrophil survival, chemotaxis, and activation, underscoring the importance of stromal amplification. Of course, our experiments do not formally exclude the possibility that proteolytically processed or post-translationally modified IL-1α could directly affect neutrophils or other myeloid cells directly.

Recently, Marteau *et al.*^63^ used single-cell and spatial transcriptomic profiling in colorectal cancer to define distinct neutrophil phenotypes and identify a neutrophil-enriched niche characterized by extensive interactions between neutrophils and fibroblasts. Their cell-cell interaction analyses indicated that fibroblast-derived CXCL1, CXCL3, and CXCL5 recruit neutrophils via CXCR1/2, and suggested that neutrophil-derived IL-1β may contribute to fibroblast polarization within these niches. Notably, the upstream signals driving the establishment of this myeloid-stromal structure remained unresolved. Our findings extend this model by identifying tumor-derived IL-1α as a key initiating signal that programs inflammatory fibroblasts and establishes spatially confined chemokine-rich stromal niches that recruit and sustain TANs. Together, these observations support a feed-forward model in which tumor cell-initiated IL-1α signaling establishes inflammatory stromal niches that may subsequently be reinforced by neutrophil-derived IL-1β, highlighting IL-1-dependent tumor-stroma–neutrophil crosstalk as a conserved organizational principle across epithelial cancers.

Building on this framework, our spatial analyses reveal that IL-1α-conditioned niches generate *CXCL8/CSF3*-co-expressing stromal zones adjacent to *IL1A*^+^ tumor islets that selectively attract TANs. Within these niches, TANs preferentially accumulate in the stromal compartment rather than tumor nests, consistent with recruitment and retention driven by stromal chemokine gradients, extracellular matrix cues, and direct stromal cell interactions. This spatial bias supports a model in which stromal cells function as the primary organizers of TAN density and positioning within the tumor, rather than neutrophils being directly recruited into tumor cell–dense regions. Importantly, this spatially confined enrichment was not observed for T cells or macrophages, highlighting a TAN-specific stromal program. Other immune populations may nonetheless be indirectly influenced by the IL-1α/iCAF/TAN axis, potentially shaping the broader tumor immune landscape. Importantly, these IL-1α-conditioned stromal niches not only determine TAN abundance but may also shape neutrophil functional state. IL-1α-activated MSCs induced a transcriptionally and phenotypically activated TAN profile characterized by enhanced survival, inflammatory marker expression (*IL1B, PD-L1, CD54*), and chemotaxis, while classical effector functions such as ROS production and degranulation remained largely unchanged. Consistently, tissue analyses revealed preservation of granule content (MPO, NE) in TANs within *IL1A*^+^ niches, whereas *IL1A*^−^ regions accumulated terminally degranulated or NETotic TANs. Together, these findings indicate that IL-1α-driven stromal niches coordinate both the spatial positioning and functional state of TANs in HNSCC.

By integrating 2D cell culture, 3D spheroids, the zebrafish xenograft model, which allows coinjection of human tumor cells and MSCs, and spatially resolved patient tissue analyses, our study links molecular signaling to tissue architecture and establishes a robust mechanistic framework for understanding neutrophil-stroma co-evolution in solid tumors. Across these models and in patient cohorts, IL-1α consistently enhanced neutrophil infiltration into tumor-MSC/stroma compartments, demonstrating the functional relevance of the stromal chemokine circuit *in vivo*. From a clinical perspective, the impact of TANs on disease progression and outcome is highly context dependent, varying with neutrophil phenotype, spatial localization, and tumor type. In many solid tumors, including HNSCC, accumulation of stromal and inflammatory TANs has been associated with immune suppression, invasion, metastasis, and poor prognosis^29,30,64^. Therapeutic strategies aimed at broadly limiting neutrophil recruitment have thus far shown limited clinical success, likely reflecting redundancy in chemokine networks and an incomplete understanding of TAN heterogeneity, spatial organization, and recruitment mechanisms^17,65^. Our data highlight IL-1α as a key mediator shaping TAN density and phenotype, suggesting that IL-1/IL-1R blockade, particularly in combination with CXCL8 blockade or immunotherapy, may represent a rational therapeutic strategy^56,66^. The *IL1A/CXCL8/CSF3* high spatial signature identified in patient tissue could serve as a biomarker to stratify patients most likely to benefit from such combinatorial approaches. In addition, previous work has shown that therapy-induced tumor cell death, such as chemotherapy, can lead to IL-1α release, which in turn can promote immune suppression and myeloid cell recruitment in the TME^67^. The specific mechanism by which therapy-induced IL-1α might program stromal fibroblasts toward an inflammatory iCAF phenotype and thereby enhance neutrophil recruitment has not been fully delineated, and our data provide novel evidence supporting this concept.

While our models and tissue analyses provide strong mechanistic insight, they cannot fully capture the complexity of the human TME, and human tumor-MSC interactions in zebrafish xenografts cannot recapitulate all immune dynamics. Neutrophil heterogeneity and potential compensatory chemokine pathways could influence outcomes. Future studies integrating *in vivo* models, spatial multiplexing, and single-cell transcriptomics will be essential to define how IL-1α interacts with other inflammatory mediators to shape TAN heterogeneity, immune suppression, and therapy response. Finally, due to the limited number of female patients in our cohort, potential sex-specific differences in IL-1α-stroma-TAN interactions could not be assessed and require validation in larger studies.

In summary, we show that IL-1α-driven tumor-stroma communication represents a central determinant of TAN-enriched niches in HNSCC, linking molecular signaling to tissue architecture and clinical prognosis. By instructing spatially confined stromal circuits, IL-1α shapes both the abundance and functional state of TANs, providing a potential target for biomarker-driven immunomodulatory strategies in neutrophil-rich tumors.

## Acknowledgements

We thank the Imaging Centre Essen (IMCES) at the Faculty of Medicine of the University of Duisburg Essen, Germany for providing access to Zeiss Axio Scan Z.1, Leica SP8 MP and IT-image analysis system. Special thanks to Anthony Squire for supporting with the spheroid migration assay analysis using Imaris 10. We thank all the blood donors and clinical personal that drew blood. We acknowledge the excellent technical assistance of Enrique de Vega Gómez and Sabine Schlink. ChatGPT (OpenAI) was used for text editing and summarizing literature. All analyses, interpretations, and conclusions remain those of the authors.

## Author Contributions

Conceptualization, EMH, SB

Methodology, EMH, KB, MS, BB, XW, AB, DMB, MR

Resources, AD, TH, StL, SB

Investigation, EMH, KB, WO, FHV, MS, BB, SiL, JAS, SR, XW, AB

Formal Analysis, EMH, KB, WO, FHV, MS, BB, SiL, JAS, XW, AB

Data Curation, EMH, KB, FHV

Writing – Original Draft, EMH, KB, SB

Writing – Review and Editing, all authors

Visualization, EMH, KB, WO, FHV, MS, BB, SiL, AB

Supervision, OS, DMB, SB

Project Administration, SB

Funding Acquisition, SB

## Methods

### Study subjects

PMNs for *in vitro* assays were isolated from the peripheral blood of healthy male and female donors. Tumor specimens and non-malignant oral cavity mucosa were collected from patients with head and neck squamous cell carcinoma during surgical resection. All procedures were conducted in accordance with the Declaration of Helsinki and approved by the local ethics committee, with written informed consent obtained from each participant. Patient characteristics are shown in Supp. Table 1a-c.

### Cell lines and generation of overexpression and knock-out models

Human carcinoma cell lines: The epidermoid carcinoma cell line A431 (ATCC CRL-1555) was cultured in DMEM with 4.5 g/L glucose (Pan-Biotech, Aidenbach, Germany), supplemented with 10% heat inactivated FCS (BioSell, Feucht, Germany) and 100 units/ml penicillin and 100 µg/mL streptomycin (Thermo Fisher Scientific, Waltham, USA). The head and neck cancer cell lines FaDu (ATCC HTB-43) and PCI13 (kindly provided by the Pittsburgh Cancer Institute) were cultured in RPMI-1640 (Pan-Biotech), supplemented with 10% FCS (BioSell) and 100 units/mL penicillin and 100 µg/mL streptomycin (Thermo Fisher Scientific, Waltham, USA). Overexpressing IL-1α FaDu cells were generated via lentiviral transduction using pLV[Exp]-eGFP-hIL1A; pLV[Exp]-eGFP-ORF served as control (both VectorBuilder ic. Chicago, USA). *CXCL8* knockout FaDu cells were generated using lentiCRISPRv2 (Addgene, Watertown, USA) with a gRNA targeting human CXCL8, followed by puromycin selection.

MSC cells were isolated from healthy oral cavity mucosa according to a previously described protocol^68^. The patientś characteristics are listed in Supp. Table 1c. The cells were immortalized between passage 1 and 3 using SV40 lentiviral particles (abm®, Richmond, BC, Canada). For imaging immortalized MSCs were lentiviral transduced with pLVX-DsRed-Monomer-N1 vector (Takara Bio Inc., Saint-Germain-en-Laye, France) and selected via puromycin. *CXCL8* knockout MSCs were generated via lentiCRISPRv2 (Addgene, Watertown, USA) carrying a gRNA targeting human CXCL8, followed by puromycin selection.

### Isolation of PBMC and PMN

Peripheral blood anticoagulated with 3.8% sodium citrate was diluted 1:1 in PBS and separated by density gradient centrifugation (Pancoll, Density 1.077 g/mL, Pan-Biotech). PBMCs were collected and thoroughly washed with PBS. PMNs were isolated by erythrocyte sedimentation using 1% polyvinyl alcohol, followed by 0.2% NaCl lysis. Isotonicity was restored with 1.2% NaCl. PBMCs and PMNs were cultured in RPMI-1640 supplemented with 10% FCS, 100 units/mL penicillin and 100 µg/mL streptomycin.

### Generation of cell supernatants and *in vitro* cell stimulation

For production of conditioned medium, cancer cells (1.5-2×10^6^ cells/mL) and MSCs (0.5×10^6^ cells/mL) were cultured for 24 h. Necrosis was induced by two freeze-thaw cycles in fresh medium with equal amounts of medium as control cells. Conditioned media were collected, centrifuged at 2000 rpm for 10 min at 4°C to remove debris, aliquoted and stored at –20°C. Tumor cells, MSCs and PMNs were stimulated with conditioned medium, as indicated.

To determine the IL-1α concentration required for CXCL8 and G-CSF induction, MSCs were treated with recombinant IL-1α at concentrations ranging from 0.01 to 10 ng/mL. Based on this titration, subsequent assays were performed with tumor-conditioned medium in the presence or absence of 500 ng/mL KINERET (anakinra), 10 µg/mL anti human IL-1α antibody, or the corresponding isotype control was added. MSCs were treated with 10 ng/mL IL-1α. For pharmacological inhibition of signaling pathways, MSCs were pre-incubated for 90 min with 10 µM NF-κB Activation Inhibitor III or 1 µM AZD1480 to block NF-κB and JAK/STAT signaling, respectively. Cells were subsequently stimulated with IL-1α in the continued presence of the inhibitors for 24 h. A detailed list of reagents is provided in Supp. Table 3.

### Cytokine / chemokine analysis

Cell culture SNs were analyzed using specific ELISAs, a human Luminex Discovery Assay (BioTechne with the Bio-Plex System (Bio-Rad, Munich, Germany), and a human Cytokine Array C5 (Ray-Biotech), following the manufacturers’ instructions. Detail of analyzed cytokines, chemokines, and corresponding assays are provided in Supp. Table 3.

### Flow cytometric analysis of stimulated MSCs

After 24 h incubation with 10 ng/mL recombinant IL-1α, MSCs were harvested and stained with fluorochrome-conjugated antibodies listed in Supp. Table 3. Stained cells were analyzed using a BD FACS Canto II flow cytometer, and date were processed with BD FACS Diva v8.01 software and FlowJo v10 (BD Bioscience).

### RNAscope analysis of stimulated MSCs

A total of 50,000 MSCs were seeded into 8-well chamber slides (Millipore). After 48 h of incubation, cells were treated with 10 ng/mL recombinant IL-1α for 6 or 24 h. Cells were washed with PBS and fixed with 4% paraformaldehyde for 30 min at room temperature. Subsequently, cells were dehydrated using a graded ethanol series, and expression of CXCL8 and CSF3 was analyzed using the RNAscope® assay for adherent cells cultured on coverslips (Advanced Cell Diagnostics [ACD], Newark, USA) according to the manufacturer’s protocol. Probes for human CXCL8 (310381-C3) and CSF3 (425301), as well as corresponding control probes, were applied. COX2 was counterstained. Nuclei were counterstained with DAPI, and cells were mounted using ProLong™ Gold Antifade Mountant (Thermo Fisher Scientific). Images were acquired using an AxioScan Z.1 microscope at 20× magnification.

### RNAseq analysis of stimulated MSCs

MSC cells were stimulated for 6 h or 24 h with 10 ng/mL IL-1α. Untreated cells were used as control. After stimulation cells were washed with PBS, and total RNA was isolated using NucleoSpin® RNA II kit (MACHEREY-NAGEL GmbH & Co. KG, Düren, Germany) according to manufacturer’s protocol. RNA concentration and integrity were assessed using a Qubit (Thermo Fisher Scientific) and an Agilent Bioanalyzer RNA nano chips (Agilent, Santa Clara, CA USA).

Library preparation was performed with Lexogens QuantSeq 3’ mRNA-Seq Library Prep Kit FWD, and sequencing was conducted on a NextSeq2000 (Illumina, San Diego, CA, USA). Sequences were trimmed with TrimGalore (v.0.6.0)^69^ and aligned with hisat2^70^. Statistical analysis was performed with R (v. 4.4.1, R Core Team (2022), R: A language and environment for statistical computing. R Foundation for Statistical Computing, Vienna, Austria. (URL https://www.R-project.org/) with the R packages DESeq2 (v 1.44)^71^, ComplexHeatmaps (v 2.20.0)^72^, umap (v 0.2.10.0) (Konopka T (2022). umap: https://cran.r-project.org/web/packages/umap/), fgsea (v 1.30) (Korotkevich G, Sukhov V, Sergushichev A. Fast gene set enrichment analysis. bioRxiv. 2019:060012) and EnhancedVolcano (v 1.22.0). Genes expressed only in one sample or with a mean expression below five were excluded from the analysis.

### Functional assays with PMNs

#### Survival assay

PMNs were cultured at 1×10^6^ cells/mL for 20 h in conditioned or control medium. 100 ng/mL G-CSF (Chugai Pharmaceutical Co., Tokyo, Japan) were used as control for apoptosis inhibition. Where indicated, 10 µg/mL of anti G-CSF or GM-CSF blocking antibodies were added (see Supp. Table 3). PMN survival was assessed by staining with Annexin V-PE and 7AAD (BD Bioscience). Stained cells were analyzed using a BD FACSCanto II flow cytometer, and data were processed using BD FACS Diva v8.01 and FlowJo v10 Software.

#### ROS assay

PMNs (1×10^6^ cells/mL) were primed for 1 h with medium, 10 ng/mL recombinant IL-1α, FaDu conditioned medium (NS or IL-1α-OE), or MSC SN pre-conditioned with FaDu NS or IL-1α-OE medium. Intracellular ROS was detected by incubation with 123-Dihydrorodamine (Thermo Fisher Scientific) for 30 min, followed by flow cytometry analysis on a BD FACS Canto II flow cytometer. Data were processed with BD FACS Diva v8.01 and FlowJo v10 Software. Extracellular ROS was measured by washing primed cells and incubating them with 100 µM luminol (Sigma-Aldrich), in the presence or absence of 12.5 nM PMA. Relative luminescence units (RLU) were recorded every 2 min over 60 min at 37°C using the Synergy 2 plate reader (BioTek Instruments, Winooski, VT, USA). The area under the curve was determined using GraphPad prism and was normalized to untreated control cells.

#### PMN degranulation, combined with Arginase assay

PMNs (1×10^6^ cells/mL) were incubated with 10 ng/mL recombinant IL-1α, MSC-conditioned, or IL-1α-stimulated MSC conditioned medium for 15, 45, and 90 min in the prescence of the Golgi inhibitor Monensin. Cells were washed, fixed, permeabilized and stained for MPO, MMP9, and Lactoferrin (see Supp. Table 3), followed by flow cytometry analysis on BD FACSCanto II. Data were processed using BD FACS Diva v8.01 and FlowJo v10 Software. To assess the release of granule proteins, SNs were collected after 90 min and analyzed for MPO, MMP9 and Arginase using ELISAs and an arginase assay kit, respectively (see Supp. Table 3).

#### Gene expression analysis/ qRT-PCR

PMNs (1×10^6^ cells/mL) were cultured for 4 h in conditioned or control medium. Total RNA was extracted using the RNeasy Mini Kit (Qiagen) according to the manufacturer’s instructions. cDNA was synthesized by reverse transcription using SuperScript™ II Reverse Transcriptase according to the manufactureŕs instructions (Thermo Fisher Scientific). Quantitative Real-time polymerase chain reaction (qRT-PCR) was performed using the Luna® Universal qPCR Master Mix (New England Biolabs) and the StepOnePlus Real-Time PCR System with StepOne software. Gene-specific primers were used to assess expression of target genes in PMNs. Sequences for forward and reverse primer are listed in Supp. Table 3. Expression levels were normalized to β-2-microglobulin (B2M) as a housekeeping gene. Relative expression was calculated using the 2ΔΔCt method.

#### PMN chemotaxis

PMNs (0.5×10^6^ cells) were allowed to migrate for 3 h through a 3 µm transwell insert (Sarstedt, Nümbrecht, Germany) towards medium or conditioned medium. Migrated cells were quantified using 123count™ eBeads (Thermo Fisher Scientific) on a BD FACS Canto II. Migration was expressed as percentage migration (migrated cells*100/ maximum migration). Where indicated 10 µg/mL of anti CXCL8 blocking antibodies were added (see Supp. Table 3).

### 3D spheroid cell culture model

Spheroids were generated from 20.000 Ds Red^+^ or non-fluorescent MSCs and 10.000 FaDu, eGFP-FaDu-NS or eGFP-FaDu-IL-1α OE cells on agarose-coated 96-well plates for 72 h. For mixed spheroids, one MSC and one FaDu spheroid were co-cultured in fresh medium for 2 h prior to the addition of 150.000 CMFDA– or DeepRed-labeled PMNs (approximate MSC:PMN ratio 1:5). After 8 h, non-migrated PMNs were removed by washing. Spheroids were fixed with 5% neutral buffered formalin, blocked overnight in 5% normal goat serum containing 0.5% Triton X-100) and counterstained with DAPI. Confocal z-stacks were acquired using a Leica TCS SP8 microscope and analyzed with LAS X, Fiji-ImageJ, and Imaris v10 software.

For tumor growth assay, tumor spheroids were cultured for 72-96 h and were subsequently treated with conditioned media from MSCs or IL-1α stimulated MSCs, as indicated. Brightfield images were acquired at 0 h and 48 h. For each spheroid, three perpendicular diameters were measured in ImageJ and the mean diameter was used for further analysis. The projected spheroid area was calculated and values at 48 h were normalized to the corresponding 0 h baseline to determine relative spheroid growth (fold change).

### Zebrafish xenograft model

FaDu cells were labeled with CellTrace Violet (5µM, C34557, Thermo Fisher Scientific) and MSCs with CellTracker Deep Red (1µM, C34565, Thermo Fisher Scientific) at 37 °C for 15 min. After washing, FaDu cells and MSCs were mixed in a ratio of 4:1 and resuspended in RPMI 1640 (Sigma-Aldrich) containing 2 mM EDTA for microinjection. Zebrafish (*Danio rerio*) embryos of the line Tg*(lyz:dsred*) were kept in E3 medium at 28.5 °C. At 2 days post fertilization (2 dpf), larvae were anesthetized in 0.016% tricaine and mounted in an agarose injection plate. A total of 600 CellTrace Violet-labeled FaDu cells and 150 CellTracker Deep Red-labeled MSCs were microinjected into the perivitelline space (PVS) in a volume of 5 nl. The injection process was performed as previously described^73^. Injected larvae were maintained in E3 medium at 34 °C. 24 h post injection (hpi), larvae were euthanized by an overdose of tricaine (0.3 mg/mL) and fixed in 4% formaldehyde solution overnight before washing for three times with PBS and storage in PBS at 4 °C. For imaging, the larvae were mounted in low-melting agarose and imaged using an upright spinning disc confocal laser microscope (Examiner, Zeiss) equipped with a confocal scanner unit CSU-X1 (Yokogawa Electric Corporation), a CCD camera (Evolve, Photometrics) and a 10×/0.3NA water immersion objective (N-Achroplan, Zeiss) and Slidebook 6.0.13 Software (3i). Neutrophil infiltration into the tumor/stroma region was quantified using ImageJ by counting DsRed-positive neutrophils within the area of FaDu cells and MSCs.

Raising and housing of adult zebrafish as well as the experiments described were performed in accordance with animal protection standards of the Ludwig-Maximilians-Universität München and approved by the government of Upper Bavaria (Regierung von Oberbayern).

### Tissue Preparation and Staining

#### Hematoxylin staining

Tumor biopsies from HNC patients were embedded in Tissue-Tek O.C.T Compound (Sakura Finetek, Umkirchen, Germany) and sectioned at 5 µm thickness. Sections were rehydrated through graded ethanol, stained with hematoxylin for 1 min and 0.5% eosin for 3 min, dehydrated, cleared in xylene and mounted using ROTI®-Histokit II (Roth, Karlsruhe Germany). Images were acquired using an AxioScan Z.1 with 20x magnification with ZEN Blue software (Carl Zeiss AG, Oberkochen, Germany).

#### Spatial gene expression analysis by RNAscope

Spatial expression of *IL1A*, *CXCL8*, and *CSF3* was analyzed using the Multiplex Fluorescent RNAscope® ISH assay (Advanced Cell Diagnostics /ACD, Newark, USA), according to the manufactureŕs protocol. Fresh frozen sections were fixed in 4% paraformaldehyde, dehydrated, and subjected to protease treatment. Probes for human *IL1A* (556791), *CXCL8* (310381-C3), *CSF3* (425301), and corresponding controls were applied. Following signal amplification and fluorophore labeling, nuclei were counterstained with DAPI and sections were mounted with ProLong™ Gold Antifade Mountant (Thermo Fisher Scientific). Images were acquired with an AxioScan Z.1 with 20x magnification and analyzed using HALO® imaging analysis platform (Indica Labs, Albuquerque, USA).

#### IF analysis of immune cell infiltrate

The spatial distribution of immune cells in HNC tissue was analyzed using a multiplex antibody panel (see Supp. Table 3). Tissue sections were sequentially incubated with uncoupled primary antibodies, flurochrome-conjugated secondary antibodies, and directly coupled primary antibodies. Nuclei were counterstained with DAPI, and sections were mounted. Imaging was performed using a Zeiss AxioScan Z.1 at 20 x magnification with ZEN Blue software. Spatial analysis was conducted using HALO® imaging platform (Indica Labs, Albuquerque, USA).

#### MACSima analysis

Multiplex immunofluorescence analysis was performed using the MACSima™ Imaging System (Miltenyi Biotech), which enables highly multiplexed cyclic imaging beyond conventional mmunofluorescence. Tissue sections were fixed with paraformaldehyde and nuclei were counterstained with DAPI. In contrast to conventional IF, the MACSima system performs sequential cycles of antibody staining, imaging, and photobleaching, allowing the detection of a large number of markers on the same tissue section (see Supp. Table 3 for antibody details). Images were stitched and pre-processed with MACS iQ View Analysis Software (Miltenyi Biotech) and further analyzed using the HALO® image analysis platform (Indica Labs, Albuquerque, USA). The phenotype of TANs was defined as CD66b^+^ and/or CD11b^+^ and/or CD15^+^. T cells were defined as CD45^+^ and CD3^+^. Macrophages were defined as CD68^+^.

### Tissue analysis

#### Tissue Classification

Tumor and stromal regions were segmented using an AI-assisted tissue classification approach based on H&E staining, tissue morphology, and intrinsic autofluorescence. For classification of MACSima images, immunofluorescent stainings of cytokeratin (tumor) and vimentin (stroma) was additionally included. Classifier performance was independently validated by two researchers to ensure accurate tissue segmentation.

#### RNA Signal Detection

RNAscope *in situ* hybridization images were analyzed to detect cytokine probe signals within tumor and stromal regions. Positive cells were identified using the FISH detection algorithm in HALO™ (Indica Labs), with signal thresholds optimized for each fluorescent channel and tumor to minimize background noise. Double-positive cells were defined as those simultaneously expressing *CXCL8* and *CSF3* above the respective thresholds. To validate RNA integrity and probe specificity, positive control probes targeting *PPIB* (*peptidylprolyl isomerase B*), *UBC*, and *POLR2A*, as well as negative control probes targeting the bacterial gene DapB (three probe set), were included according to the manufacturer’s protocol. As expected, no signal was detected in the 488 channel in negative control slides. Minimal background signal was observed in the 555 and 647 channels, likely due to intrinsic tissue autofluorescence or nonspecific hybridization; therefore, channel thresholds were conservatively optimized on a per-sample basis. Sample and staining validation were independently performed by two researchers.

#### Spatial zonal analysis

Tumor regions with or without *IL1A* expression were manually annotated based on a density heatmaps, extending up to the tumor-stroma boundary. Tumor areas lacking contact with surrounding stroma were excluded from the analysis. Using the HALO® image analysis platform, concentric stromal zones were generated extending 200 µm outward from the annotated tumor boundary, with each zone measuring 50 µm in width. The stroma annotation generated by the tissue classifier was used to exclude tumor tissue from all zones, ensuring that analysis was restricted to stromal regions only. Within each stromal zone, *CXCL8^+^/CSF3*^+^ double positive stromal cells were quantified and normalized by stromal area (cells/mm²). These data were then used to assess spatial gradients in cytokine expression relative to proximity to the *IL1A*-negative and *IL1A*-positive tumor regions.

### TCGA Transcriptomic Analysis

Transcriptomic data from 515 tumor and 44 matched tumor-adjacent normal HNSCC samples available in The Cancer Genome Atlas (TCGA) were accessed through the cBioPortal platform (https://www.cbioportal.org/). Transcriptomic data analyses were conducted using R (v 4.5.1; R Foundation for Statistical Computing, Vienna, Austria, https://www.R-project.org/)) in RStudio (v 2025.05.1). Data were normalized to z-scores using *zscore* function from the package *liver* (v 1.26). MSCs signature was calculated as the mean value of the top 100 genes significantly upregulated in MSCs stimulated with IL-1α for 24h, compared to untreated cells (Supp. Table 2). TAN signatures were derived from four independent studies^41–44^, and calculated as the mean value of the genes described in each study. Spearman correlations were calculated using *rcorr* function and plots were generated using the function *corrplot*, both from package *Hmisc* (v 5.2.4).

### Statistical analysis

Statistical analyses were performed using GraphPad Prism 10. Data were analyzed using paired or unpaired two-tailed *t* tests, one-way ANOVA followed by Tukey’s multiple-comparisons test, or Wilcoxon matched-pairs signed-rank test, as indicated in the figure legends. Data are presented as mean ± SD or mean ± SEM, as specified. A p value < 0.05 was considered for statistical relevance.

## Supplements

### Supplementary Figures

**Supplementary Figure 1:**
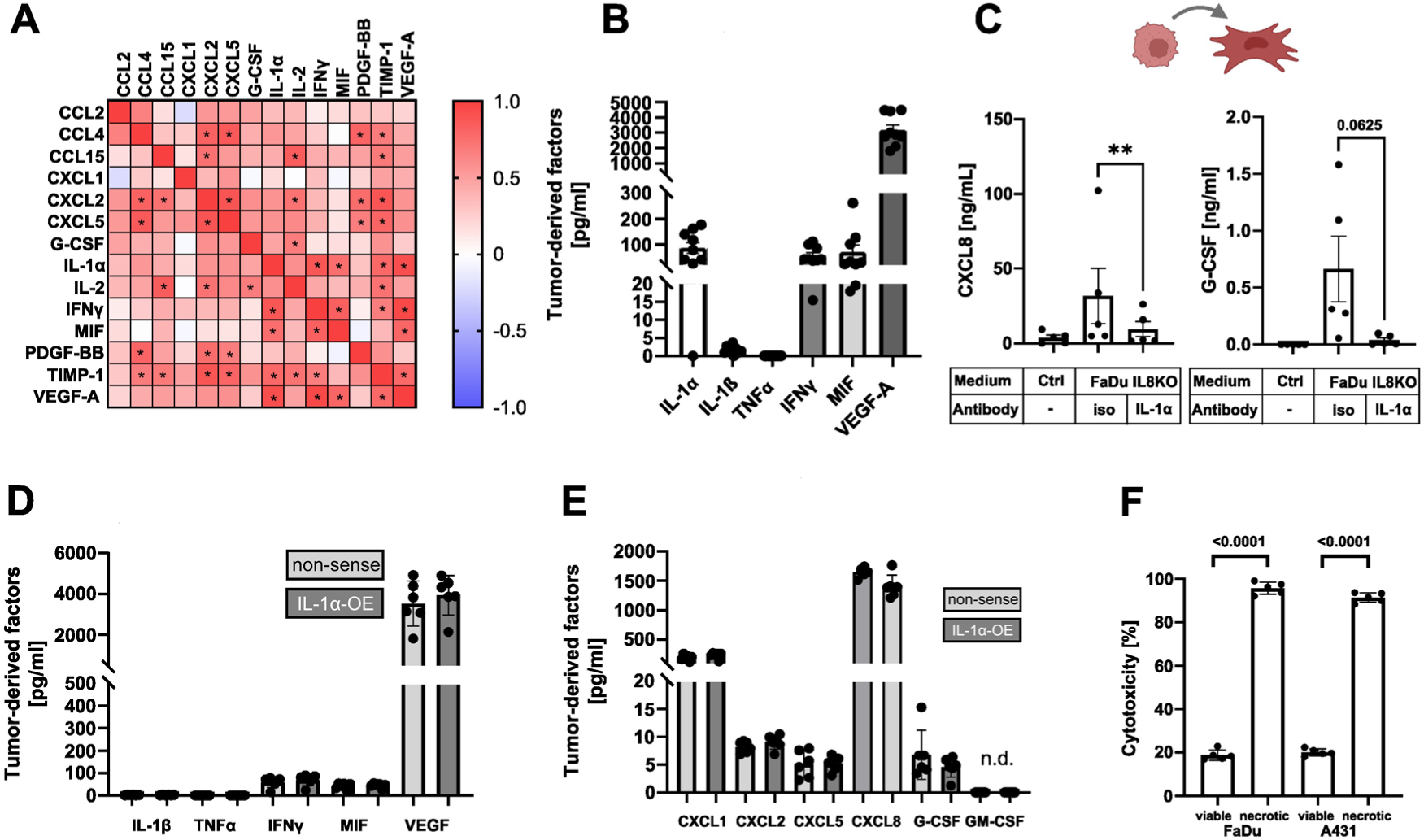
Tumor-derived IL-1α induces CXCL8 and G-CSF release by stroma cells. (A,B) Tumor-derived cytokines were quantified by Luminex analysis. (A) Data are displayed as Pearsońs correlation matrix. Significant correlations among tumor-derived factors are indicated by black asterisks (B). Quantification of tumor-derived mediators known to be involved in tumor-stroma communication. (C) MSCs were treated with FaDu-CXCL8KO tumor conditioned medium in the presence of 10 µg/mL IL-1α neutralizing antibody or isotype control. Released CXCL8 and G-CSF were quantified by ELISA. (D) Quantification of tumor-derived mediators of tumor-stroma communication in non-sense and IL-1α overexpressing (IL-1α-OE) cells was performed by Luminex. (E) Quantification of tumor-derived factors implicated in neutrophil recruitment and survival in SN from non-sense and IL-1α-OE cells (Luminex and CXCL8-ELISA). (F) LDH assay quantifying cytotoxicity in SN of viable and necrotic FaDu and A431 tumor cells. Statistical analysis was performed using paired *t*-tests (C,F) and unpaired *t*-tests (D). Data are presented as mean ± SEM (C) and mean ± SD (D,E), p values are indicated.

**Supplementary Figure 2:**
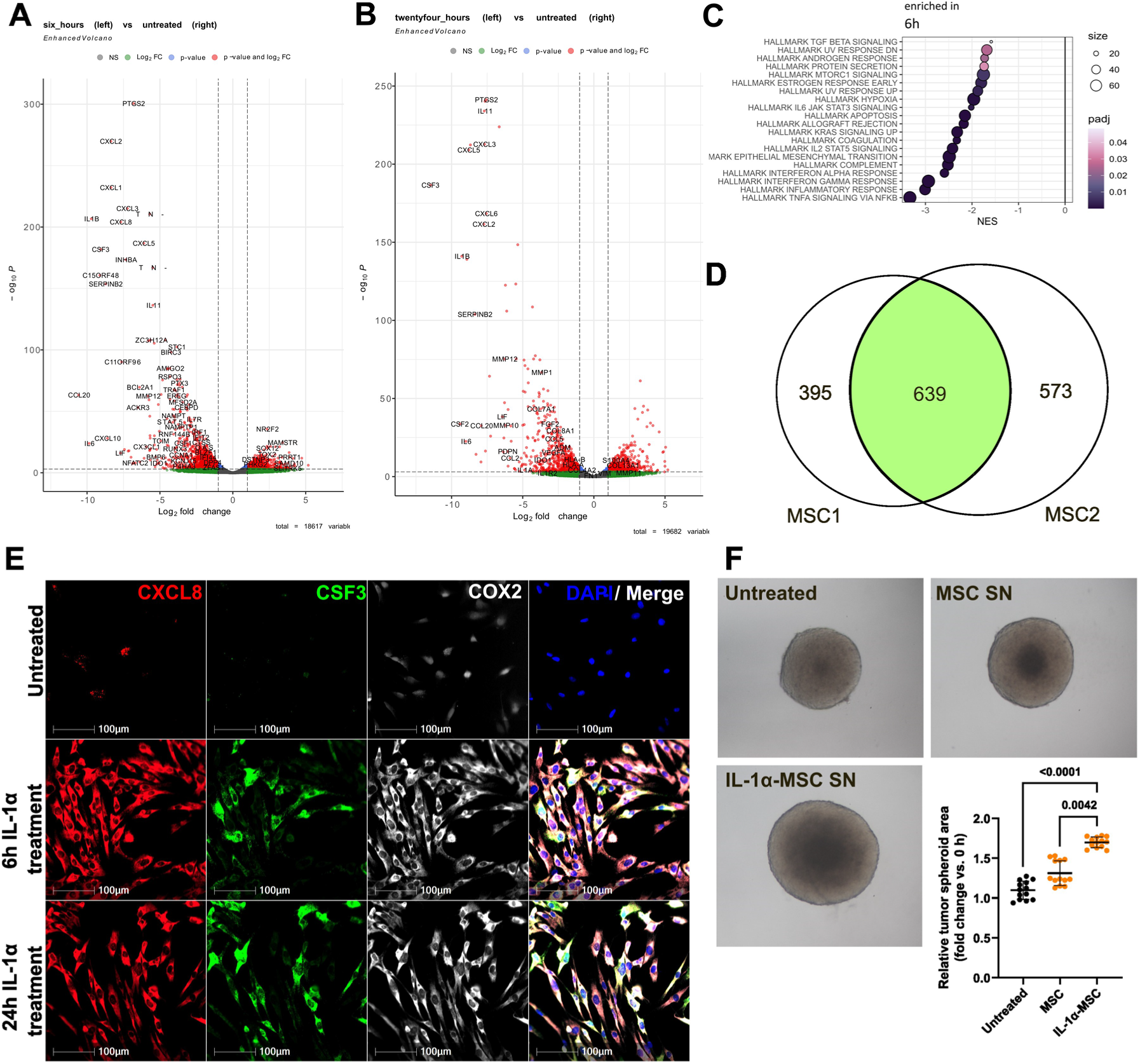
IL-1α treatment induces an inflammatory CAF-phenotype in MSCs. MSCs were treated with recombinant IL-1α for 6 and 24 h. Gene expression was analyzed by Bulk RNAseq. Volcano plots are shown for untreated vs. 6 h (A), and untreated vs. 24 h IL-1α treatment (B). (C) Enriched hallmark pathways are shown for untreated vs. 6 h. (D) Venn diagram showing the number of upregulated DEGs in two MSC cell lines after 6 h of IL-1α treatment. (E) MSC cells were treated for 6 h and 24 h with IL-1α and were analyzed by RNAscope for the expression of CXCL8 (red) and CSF3 (green) with consecutive COX2 immunostaining (white). (F) FaDu spheroids were cultured in the presence or absence of SNs derived from MSCs or IL-1α-stimulated MSCs. Relative tumor spheroid area was quantified for two independent experiments. Statistical analysis was performed using Mann-Whitney test. Data depict mean ± SD. *P*-values are indicated.

**Supplementary Figure 3:**
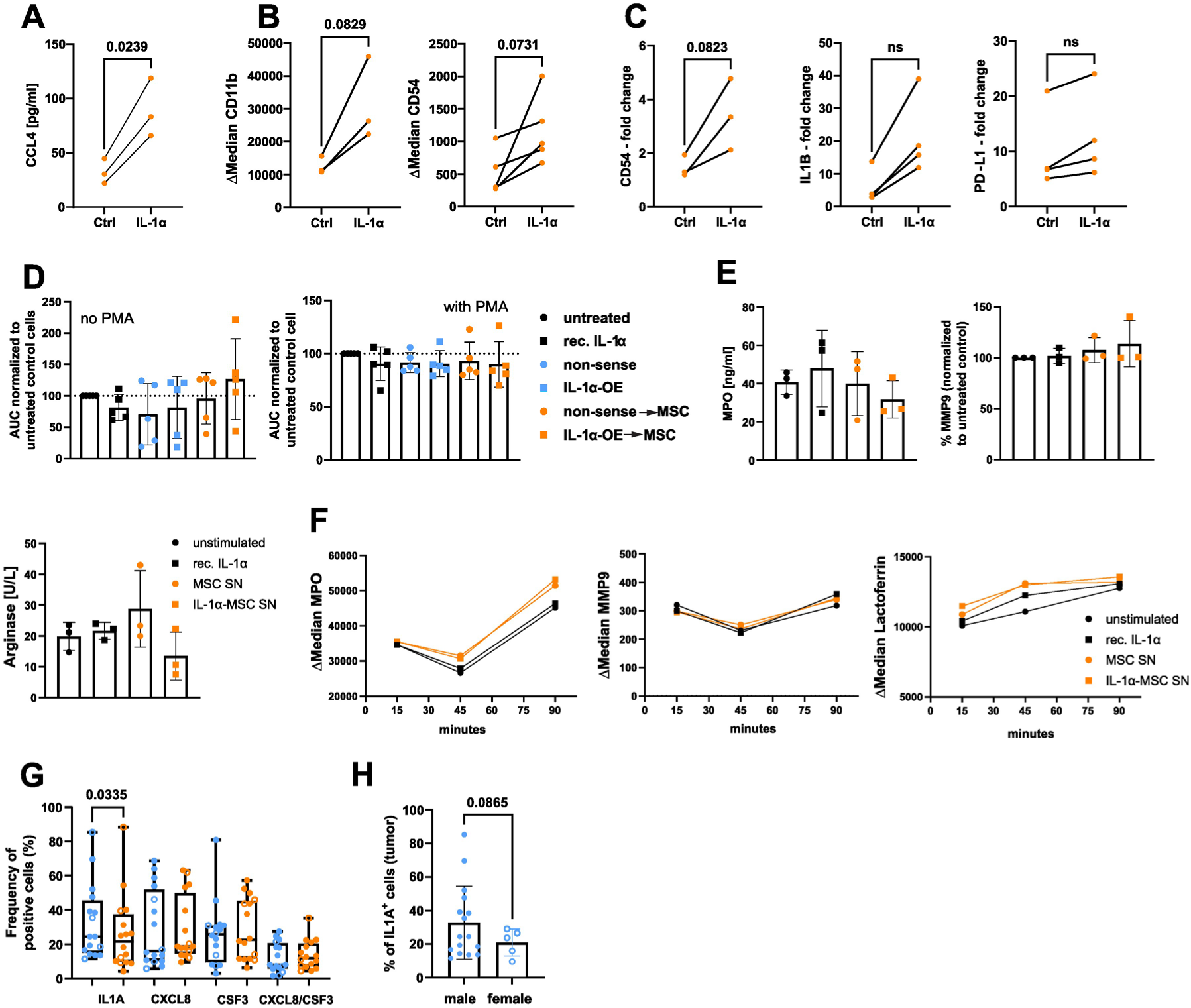
MSC-derived factors modulate PMN activation. PMNs were cultured in SN derived from untreated MSC control (Ctrl) or IL-1α-treated MSC cells. (A) CCL4 release by PMNs was measured by ELISA. (B) Surface activation markers (CD11b, CD54) on PMNs were assessed by flow cytometry. (C) CD54, IL1B, and PD-L1 transcript levels were analyzed by qRT-PCR. (D) Intracellular ROS production was quantified by luminol assay in the presence or absence of recombinant IL-1α, and tumor or tumor-primed MSC SNs. ROS was measured both with and without PMA (left: without PMA, right: with PMA). (E) PMNs were cultured with or without recombinant IL-1α and MSC or IL-1α-treated MSC SNs. Release of MPO, MMP9, and Arginase was analyzed by ELISA and arginase activity assay. (F) Degranulation of PMNs was measured by flow cytometry, data are presented as mean, n=3. G) Frequencies of cytokine positive cells in tumor (blue) and stroma (orange) across patients (n = 21), analyzed by RNAscope. H) Frequency of IL1A⁺ tumor cells stratified by sex. Statistical analysis was performed using the Wilcoxon matched-pairs signed-rank test (A–D, G,H). Data are presented as before-after (A-C), mean ± SD (D, E, H), and box-and-whiskers (G; min to max). *P* values are indicated.

### Supplementary Tables

**Supplementary Table 1a:** Baseline clinicopathological characteristics of cohort 1 (Fig. 1)

|  | n | % |
| --- | --- | --- |
| **ALL samples** | 83 | 100 |
| **Sex** |  |  |
| Male | 53 | 63,9 |
| Female | 30 | 36,1 |
| **mean age** |  |  |
| Male | 61,0+/-10,1 |  |
| Female | 63,0+/-7,5 |  |
| **Tumor localization** |  |  |
| Oropharynx | 80 | 96,4 |
| Oral cavity | 3 | 3,6 |
| **T category** |  |  |
| T1 | 4 | 4,8 |
| T2 | 29 | 34,9 |
| T3 | 31 | 37,3 |
| T4 | 20 | 24,1 |
| **Lymph node involvement** |  |  |
| N0 | 18 | 21,7 |
| N1 | 17 | 20,5 |
| N2abc | 47 | 56,6 |
| N3 | 2 | 2,4 |
| **Distant metastasis** |  |  |
| M0 | 0 | 98,8 |
| M1 | 1 | 1,2 |
| **HPV Status of OSCC** |  |  |
| p16+ | 27 | 33,8 |
| p16- | 32 | 40 |
| unkown | 21 | 26,3 |

**Supplementary Table 1b:** Baseline clinicopathological characteristics of cohort 2 (Fig. 7,8, Supp Fig. 4)

|  | n | % |
| --- | --- | --- |
| **ALL samples** | 21 | 100 |
| **Sex** |  |  |
| Male | 16 | 76,2 |
| Female | 5 | 23,8 |
| **mean age** |  |  |
| Male | 64,93 +/-10,3 |  |
| Female | 67,6 +/- 7,5 |  |
| **Tumor localization** |  |  |
| Oropharynx | 16 | 76,2 |
| Oral cavity | 5 | 23,8 |

| T category |  |  |
| --- | --- | --- |
| T1 | 1 | 4,8 |
| T2 | 9 | 42,9 |
| T3 | 4 | 19 |
| T4 | 7 | 33,3 |
| Lymph node involvement |  |  |
| N0 | 8 | 38,1 |
| N1 | 4 | 19 |
| N2abc | 7 | 33,3 |
| N3 | 2 | 9,5 |
| Distant metastasis |  |  |
| M0 | 20 | 95,2 |
| M1 | 1 | 4,8 |
| HPV Status of OSCC |  |  |
| p16+ | 9 | 56,3 |
| p16- | 6 | 37,5 |
| other | 1 | 6,3 |

**Supplementary Table 1c:** Baseline clinicopathological characteristics of patient-derived MSCs.

|  | n | % |
| --- | --- | --- |
| **ALL samples** | 12 | 100 |
| Sex |  |  |
| Male | 9 | 75 |
| Female | 3 | 25 |
| mean age |  |  |
| Male | 61,67 +/-6,4 |  |
| Female | 65,55 +/- 10,5 |  |
| Tumor localization |  |  |
| Oropharynx | 5 | 42 |
| Oral cavity | 3 | 25 |
| Larynx | 3 | 25 |
| Hypopharynx | 1 | 8 |
| T category |  |  |
| T1 | 2 | 17 |
| T2 | 7 | 58 |
| T3 | 3 | 25 |
| T4 | 1 | 8 |
| Lymph node involvement |  |  |
| N0 | 7 | 58 |
| N1 | 0 | 0 |
| N2abc | 3 | 25 |
| N3 | 2 | 17 |
| Distant metastasis |  |  |
| M0 | 12 | 100 |
| M1 | 0 | 0 |

**Supplementary Table 2.** (provided as separate Excel file)

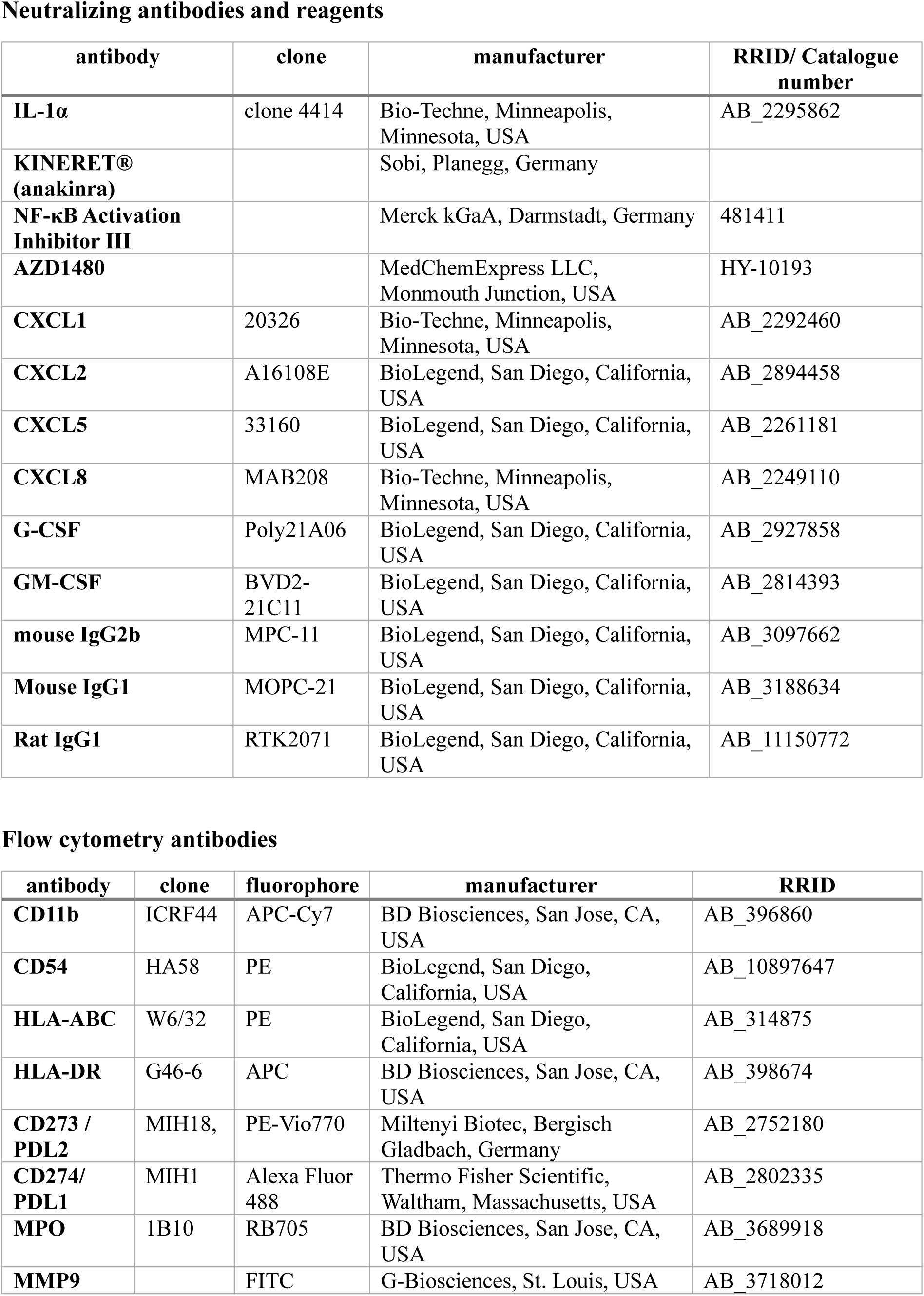
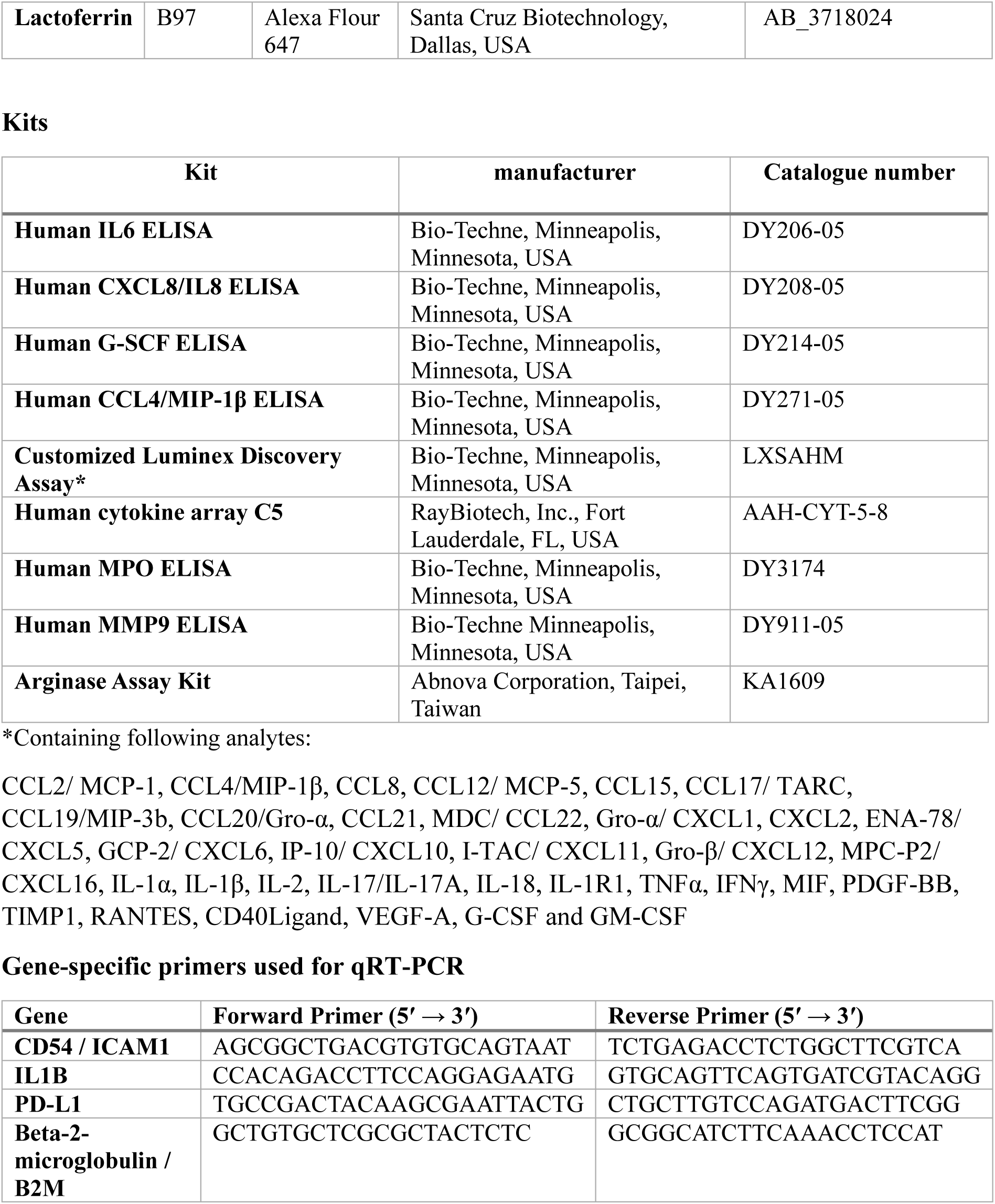
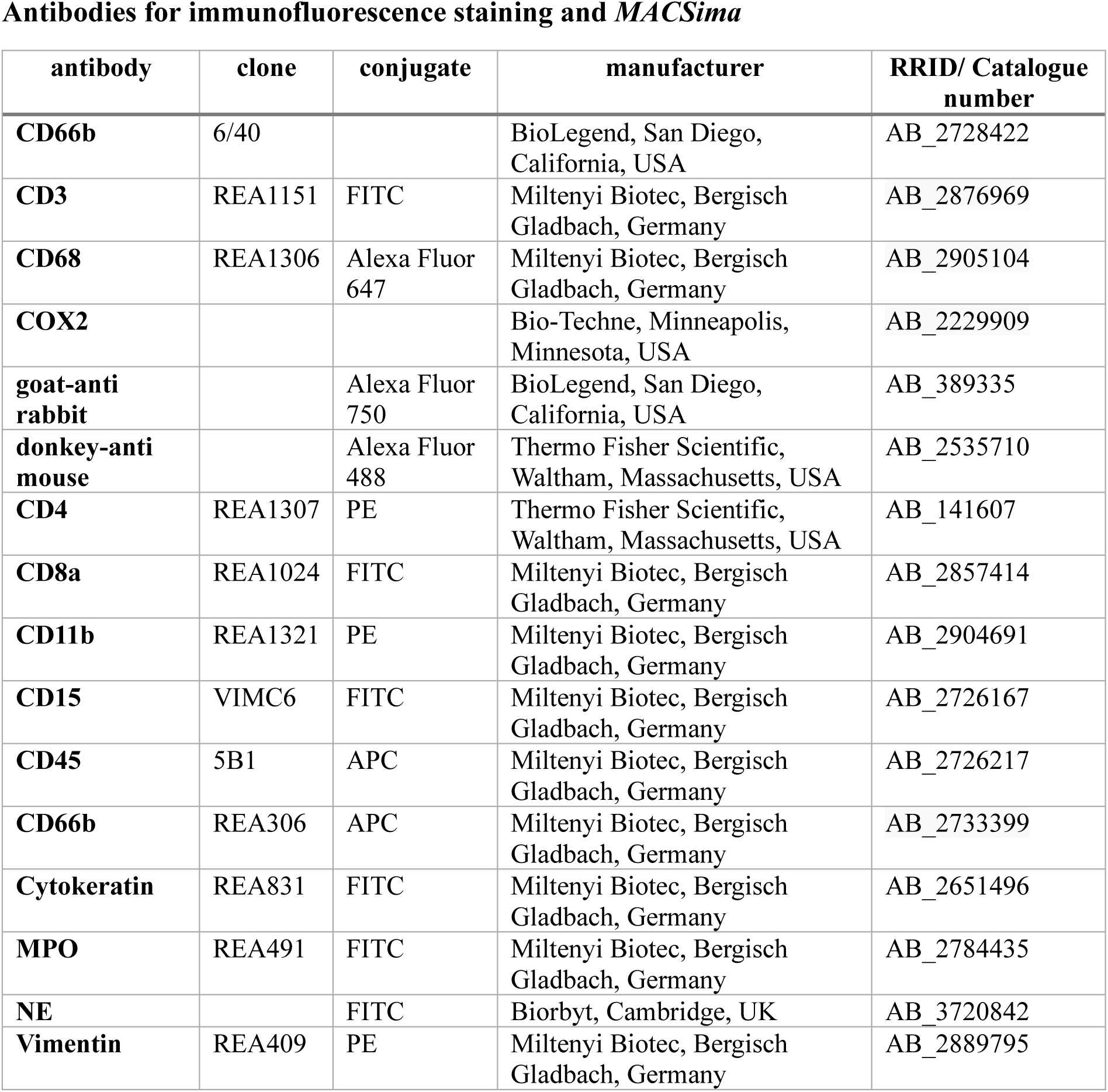
Supplementary Table 3.

## References

1 Li, Q., Tie, Y., Alu, A., Ma, X. & Shi, H. Targeted therapy for head and neck cancer: signaling pathways and clinical studies. Signal Transduct Target Ther 8, 31 (2023). 10.1038/s41392-022-01297-0

2 Canning, M. et al. Heterogeneity of the Head and Neck Squamous Cell Carcinoma Immune Landscape and Its Impact on Immunotherapy. Front Cell Dev Biol 7, 52 (2019). 10.3389/fcell.2019.00052

3 Rad, H. S. et al. Understanding the tumor microenvironment in head and neck squamous cell carcinoma. Clin Transl Immunology 11, e1397 (2022). 10.1002/cti2.1397

4 Karunasagara, S. et al. Tissue Mechanics and Hedgehog Signaling Crosstalk as a Key Epithelial-Stromal Interplay in Cancer Development. Adv Sci (Weinh) 11, e2400063 (2024). 10.1002/advs.202400063

5 Sahai, E. et al. A framework for advancing our understanding of cancer-associated fibroblasts. Nat Rev Cancer 20, 174–186 (2020). 10.1038/s41568-019-0238-1

6 Chen, Y., McAndrews, K. M. & Kalluri, R. Clinical and therapeutic relevance of cancer-associated fibroblasts. Nat Rev Clin Oncol 18, 792–804 (2021). 10.1038/s41571-021-00546-5

7 Koncina, E. et al. IL1R1(+) cancer-associated fibroblasts drive tumor development and immunosuppression in colorectal cancer. Nature communications 14, 4251 (2023). 10.1038/s41467-023-39953-w

8 Biffi, G. et al. IL1-Induced JAK/STAT Signaling Is Antagonized by TGFbeta to Shape CAF Heterogeneity in Pancreatic Ductal Adenocarcinoma. Cancer Discov 9, 282–301 (2019). 10.1158/2159-8290.CD-18-0710

9 Dominguez, C. X. et al. Single-Cell RNA Sequencing Reveals Stromal Evolution into LRRC15(+) Myofibroblasts as a Determinant of Patient Response to Cancer Immunotherapy. Cancer Discov 10, 232–253 (2020). 10.1158/2159-8290.Cd-19-0644

10 Quail, D. F. & Joyce, J. A. Microenvironmental regulation of tumor progression and metastasis. Nat Med 19, 1423–1437 (2013). 10.1038/nm.3394

11 Joyce, J. A. & Pollard, J. W. Microenvironmental regulation of metastasis. Nat Rev Cancer 9, 239–252 (2009). 10.1038/nrc2618

12 Kalluri, R. & Zeisberg, M. Fibroblasts in cancer. Nat Rev Cancer 6, 392–401 (2006). 10.1038/nrc1877

13 Hanahan, D. & Weinberg, R. A. Hallmarks of cancer: the next generation. Cell 144, 646–674 (2011). 10.1016/j.cell.2011.02.013

14 Trellakis, S. et al. Peripheral blood neutrophil granulocytes from patients with head and neck squamous cell carcinoma functionally differ from their counterparts in healthy donors. Int. J. Immunopathol. Pharmacol 24, 683–693 (2011). 10.1177/039463201102400314

15 Si, Y. et al. Multidimensional Imaging Provides Evidence for Down-Regulation of T cell Effector Function by MDSC in Human Cancer Tissue. Science immunology 4 (2019). 10.1126/sciimmunol.aaw9159

16 SenGupta, S., Hein, L. E. & Parent, C. A. The Recruitment of Neutrophils to the Tumor Microenvironment Is Regulated by Multiple Mediators. Front Immunol 12, 734188 (2021). 10.3389/fimmu.2021.734188

17 Powell, D. R. & Huttenlocher, A. Neutrophils in the Tumor Microenvironment. Trends Immunol 37, 41–52 (2016). 10.1016/j.it.2015.11.008

18 Ghodsi, A., Hidalgo, A. & Libreros, S. Lipid mediators in neutrophil biology: inflammation, resolution and beyond. Curr Opin Hematol 31, 175–192 (2024). 10.1097/moh.0000000000000822

19 McFarlane, A. J., Fercoq, F., Coffelt, S. B. & Carlin, L. M. Neutrophil dynamics in the tumor microenvironment. J Clin Invest 131 (2021). 10.1172/jci143759

20 Zarbock, A. & Ley, K. Neutrophil adhesion and activation under flow. Microcirculation 16, 31–42 (2009). 10.1080/10739680802350104

21 Smith, C. W. & Anderson, D. C. PMN adhesion and extravasation as a paradigm for tumor cell dissemination. Cancer Metastasis Rev 10, 61–78 (1991). 10.1007/bf00046844

22 Zarbock, A. & Ley, K. Mechanisms and consequences of neutrophil interaction with the endothelium. Am J Pathol 172, 1–7 (2008). 10.2353/ajpath.2008.070502

23 Höing, B. et al. Stromal versus tumoral inflammation differentially contribute to metastasis and poor survival in laryngeal squamous cell carcinoma. Oncotarget 9, 8415–8426 (2018). 10.18632/oncotarget.23865

24 Sody, S. et al. Distinct Spatio-Temporal Dynamics of Tumor-Associated Neutrophils in Small Tumor Lesions. Front Immunol 10, 1419 (2019). 10.3389/fimmu.2019.01419

25 Papayannopoulos, V., Metzler, K. D., Hakkim, A. & Zychlinsky, A. Neutrophil elastase and myeloperoxidase regulate the formation of neutrophil extracellular traps. J Cell Biol 191, 677–691 (2010). 10.1083/jcb.201006052

26 Burn, G. L. et al. Myeloperoxidase transforms chromatin into neutrophil extracellular traps. Nature 647, 747–756 (2025). 10.1038/s41586-025-09523-9

27 Antuamwine, B. B. et al. N1 versus N2 and PMN-MDSC: A critical appraisal of current concepts on tumor-associated neutrophils and new directions for human oncology. Immunol Rev 314, 250–279 (2023). 10.1111/imr.13176

28 Quail, D. F. et al. Neutrophil phenotypes and functions in cancer: A consensus statement. The Journal of experimental medicine 219 (2022). 10.1084/jem.20220011

29 Coffelt, S. B., Wellenstein, M. D. & de Visser, K. E. Neutrophils in cancer: neutral no more. Nat Rev Cancer 16, 431–446 (2016). 10.1038/nrc.2016.52

30 Shaul, M. E. & Fridlender, Z. G. Tumour-associated neutrophils in patients with cancer. Nat Rev Clin Oncol 16, 601–620 (2019). 10.1038/s41571-019-0222-4

31 Hansen, N. et al. Blocking IL1RAP on cancer-associated fibroblasts in pancreatic ductal adenocarcinoma suppresses IL-1-induced neutrophil recruitment. J Immunother Cancer 12 (2024). 10.1136/jitc-2024-009523

32 Tjomsland, V. et al. Interleukin 1α sustains the expression of inflammatory factors in human pancreatic cancer microenvironment by targeting cancer-associated fibroblasts. Neoplasia (New York, N.Y.) 13, 664–675 (2011). 10.1593/neo.11332

33 Tran, L. L., Dang, T., Thomas, R. & Rowley, D. R. ELF3 mediates IL-1α induced differentiation of mesenchymal stem cells to inflammatory iCAFs. Stem Cells 39, 1766–1777 (2021). 10.1002/stem.3455

34 Elyada, E. et al. Cross-Species Single-Cell Analysis of Pancreatic Ductal Adenocarcinoma Reveals Antigen-Presenting Cancer-Associated Fibroblasts. Cancer Discov 9, 1102–1123 (2019). 10.1158/2159-8290.CD-19-0094

35 Hu, B. et al. Subpopulations of cancer-associated fibroblasts link the prognosis and metabolic features of pancreatic ductal adenocarcinoma. Ann Transl Med 10, 262 (2022). 10.21037/atm-22-407

36 Petrov, P. B., Considine, J. M., Izzi, V. & Naba, A. Matrisome AnalyzeR – a suite of tools to annotate and quantify ECM molecules in big datasets across organisms. J Cell Sci 136 (2023). 10.1242/jcs.261255

37 Kennel, K. B., Bozlar, M., De Valk, A. F. & Greten, F. R. Cancer-Associated Fibroblasts in Inflammation and Antitumor Immunity. Clinical cancer research: an official journal of the American Association for Cancer Research 29, 1009–1016 (2023). 10.1158/1078-0432.Ccr-22-1031

38 Zhang, Q. et al. Integrated analysis of single-cell RNA-seq and bulk RNA-seq reveals distinct cancer-associated fibroblasts in head and neck squamous cell carcinoma. Ann Transl Med 9, 1017 (2021). 10.21037/atm-21-2767

39 Obradovic, A. et al. Immunostimulatory Cancer-Associated Fibroblast Subpopulations Can Predict Immunotherapy Response in Head and Neck Cancer. Clinical cancer research: an official journal of the American Association for Cancer Research 28, 2094–2109 (2022). 10.1158/1078-0432.CCR-21-3570

40 Choi, J. H. et al. Single-cell transcriptome profiling of the stepwise progression of head and neck cancer. Nature communications 14, 1055 (2023). 10.1038/s41467-023-36691-x

41 Ng, M., Cerezo-Wallis, D., Ng, L. G. & Hidalgo, A. Adaptations of neutrophils in cancer. Immunity 58, 40–58 (2025). 10.1016/j.immuni.2024.12.009

42 Wu, Y. et al. Neutrophil profiling illuminates anti-tumor antigen-presenting potency. Cell 187, 1422–1439.e1424 (2024). 10.1016/j.cell.2024.02.005

43 Cerezo-Wallis, D. et al. Architecture of the neutrophil compartment. Nature (2025). 10.1038/s41586-025-09807-0

44 Zilionis, R. et al. Single-Cell Transcriptomics of Human and Mouse Lung Cancers Reveals Conserved Myeloid Populations across Individuals and Species. Immunity 50, 1317–1334.e1310 (2019). 10.1016/j.immuni.2019.03.009

45 Campbell, J. D. et al. Genomic, Pathway Network, and Immunologic Features Distinguishing Squamous Carcinomas. Cell reports 23, 194–212.e196 (2018). 10.1016/j.celrep.2018.03.063

46 Cerami, E. et al. The cBio cancer genomics portal: an open platform for exploring multidimensional cancer genomics data. Cancer Discov 2, 401–404 (2012). 10.1158/2159-8290.Cd-12-0095

47 Gao, J. et al. Integrative analysis of complex cancer genomics and clinical profiles using the cBioPortal. Sci Signal 6, pl1 (2013). 10.1126/scisignal.2004088

48 de Bruijn, I. et al. Analysis and Visualization of Longitudinal Genomic and Clinical Data from the AACR Project GENIE Biopharma Collaborative in cBioPortal. Cancer research 83, 3861–3867 (2023). 10.1158/0008-5472.Can-23-0816

49 Wang, W. et al. Development and Validation of a Pathomics Model Using Machine Learning to Predict CXCL8 Expression and Prognosis in Head and Neck Cancer. Clin Exp Otorhinolaryngol 17, 85–97 (2024). 10.21053/ceo.2023.00026

50 Zhao, Z. et al. Expression of chemokine CXCL8/9/10/11/13 and its prognostic significance in head and neck cancer. Medicine (Baltimore) 101, e29378 (2022). 10.1097/md.0000000000029378

51 Hamilton, J. A. GM-CSF in inflammation. The Journal of experimental medicine 217 (2020). 10.1084/jem.20190945

52 Kasahara, S. et al. Role of Granulocyte-Macrophage Colony-Stimulating Factor Signaling in Regulating Neutrophil Antifungal Activity and the Oxidative Burst During Respiratory Fungal Challenge. J Infect Dis 213, 1289–1298 (2016). 10.1093/infdis/jiw054

53 Zhang, W., Borcherding, N. & Kolb, R. IL-1 Signaling in Tumor Microenvironment. Adv Exp Med Biol 1240, 1–23 (2020). 10.1007/978-3-030-38315-2_1

54 Di Paolo, N. C. & Shayakhmetov, D. M. Interleukin 1α and the inflammatory process. Nature immunology 17, 906–913 (2016). 10.1038/ni.3503

55 Lavie, D., Ben-Shmuel, A., Erez, N. & Scherz-Shouval, R. Cancer-associated fibroblasts in the single-cell era. Nat Cancer 3, 793–807 (2022). 10.1038/s43018-022-00411-z

56 Dinarello, C. A. Overview of the IL-1 family in innate inflammation and acquired immunity. Immunol Rev 281, 8–27 (2018). 10.1111/imr.12621

57 Rider, P. et al. IL-1α and IL-1β recruit different myeloid cells and promote different stages of sterile inflammation. J Immunol 187, 4835–4843 (2011). 10.4049/jimmunol.1102048

58 Carmi, Y. et al. The role of IL-1β in the early tumor cell-induced angiogenic response. J Immunol 190, 3500–3509 (2013). 10.4049/jimmunol.1202769

59 Voronov, E. et al. IL-1 is required for tumor invasiveness and angiogenesis. Proc Natl Acad Sci U S A 100, 2645–2650 (2003). 10.1073/pnas.0437939100

60 Figari, I. S., Mori, N. A. & Palladino, M. A., Jr. Regulation of neutrophil migration and superoxide production by recombinant tumor necrosis factors-alpha and –beta: comparison to recombinant interferon-gamma and interleukin-1 alpha. Blood 70, 979–984 (1987).

61 Sullivan, G. W., Carper, H. T., Sullivan, J. A., Murata, T. & Mandell, G. L. Both recombinant interleukin-1 (beta) and purified human monocyte interleukin-1 prime human neutrophils for increased oxidative activity and promote neutrophil spreading. J Leukoc Biol 45, 389–395 (1989). 10.1002/jlb.45.5.389

62 Kharazmi, A., Nielsen, H. & Bendtzen, K. Recombinant interleukin 1 alpha and beta prime human monocyte superoxide production but have no effect on chemotaxis and oxidative burst response of neutrophils. Immunobiology 177, 32–39 (1988). 10.1016/s0171-2985(88)80089-5

63 Marteau, V. et al. Single-cell integration and multi-modal profiling reveals phenotypes and spatial organization of neutrophils in colorectal cancer. Cancer Cell 44, 146–165.e114 (2026). 10.1016/j.ccell.2025.12.003

64 Jaillon, S. et al. Neutrophil diversity and plasticity in tumour progression and therapy. Nat Rev Cancer 20, 485–503 (2020). 10.1038/s41568-020-0281-y

65 Highfill, S. L. et al. Disruption of CXCR2-mediated MDSC tumor trafficking enhances anti-PD1 efficacy. Sci Transl Med 6, 237ra267 (2014). 10.1126/scitranslmed.3007974

66 Xiao, Z., Singh, S. & Singh, M. Improving cancer immunotherapy by targeting IL-1. Oncoimmunology 10, 2008111 (2021). 10.1080/2162402x.2021.2008111

67 Hänggi, K. et al. Interleukin-1α release during necrotic-like cell death generates myeloid-driven immunosuppression that restricts anti-tumor immunity. Cancer Cell 42, 2015–2031.e2011 (2024). 10.1016/j.ccell.2024.10.014

68 Schuler, P. J. & Brandau, S. Adenosine Producing Mesenchymal Stem Cells. Stem Cells 35, 1647–1648 (2017). 10.1002/stem.2532

69 Kechin, A., Boyarskikh, U., Kel, A. & Filipenko, M. cutPrimers: A New Tool for Accurate Cutting of Primers from Reads of Targeted Next Generation Sequencing. J Comput Biol 24, 1138–1143 (2017). 10.1089/cmb.2017.0096

70 Kim, D., Paggi, J. M., Park, C., Bennett, C. & Salzberg, S. L. Graph-based genome alignment and genotyping with HISAT2 and HISAT-genotype. Nat Biotechnol 37, 907–915 (2019). 10.1038/s41587-019-0201-4

71 Love, M. I., Huber, W. & Anders, S. Moderated estimation of fold change and dispersion for RNA-seq data with DESeq2. Genome Biol 15, 550 (2014). 10.1186/s13059-014-0550-8

72 Gu, Z., Eils, R. & Schlesner, M. Complex heatmaps reveal patterns and correlations in multidimensional genomic data. Bioinformatics 32, 2847–2849 (2016). 10.1093/bioinformatics/btw313

73 Martinez-Lopez, M., Póvoa, V. & Fior, R. Generation of Zebrafish Larval Xenografts and Tumor Behavior Analysis. J Vis Exp (2021). 10.3791/62373

